# Selection on network dynamics drives differential rates of protein domain evolution

**DOI:** 10.1101/026658

**Authors:** Brian K. Mannakee, Ryan N. Gutenkunst

## Abstract

The long-held principle that functionally important proteins evolve slowly has recently been challenged by studies in mice and yeast showing that the severity of a protein knockout only weakly predicts that protein’s rate of evolution. However, the relevance of these studies to evolutionary changes within proteins is unknown, because amino acid substitutions, unlike knockouts, often only slightly perturb protein activity. To quantify the phenotypic effect of small biochemical perturbations, we developed an approach to use computational systems biology models to measure the influence of individual reaction rate constants on network dynamics. We show that this dynamical influence is predictive of protein domain evolutionary rate in vertebrates and yeast, even after controlling for expression level and breadth, network topology, and knockout effect. Thus, our results not only demonstrate the importance of protein domain function in determining evolutionary rate, but also the power of systems biology modeling to uncover unanticipated evolutionary forces.

---

Over evolutionary time, every protein accumulates amino acid changes at its own characteristic rate, which Zuckerkandl and Pauling likened to the ticking of a molecular clock [1]. Remarkably, this evolutionary rate varies by orders of magnitude among proteins. Understanding the determinants of this variation is a fundamental goal in molecular evolution research [2, 3, 4, 5]. Early theoretical work suggested that functional constraints within proteins [1] and the functional importance of each protein to the organism [6, 7] would be key factors in determining evolutionary rates. Yet, empirical studies using knockouts have observed only weak effects. In bacteria [8, 9], yeast [10, 11], and mammals [12] knockout studies conclude that essential proteins evolve only slightly more slowly than non-essential proteins. Moreover, among non-essential genes in yeast, there is little to no correlation between the effect of a protein knockout on growth rate, in a wide range of conditions, and that protein’s evolutionary rate [13, 14, 11], particularly when controlling for expression level [15]. This poor correlation between knockout effects and rates of protein evolution has led some researchers to conclude that function-specific selection plays little role in determining evolutionary rates [4, 5]. This conclusion is, however, contrary to theoretical expectations, the intuition of most molecular biologists, and the reasoning behind much of comparative genomics [16], motivating our search for an alternative measure of protein function.

We reasoned that knockouts do not mimic evolutionarily relevant mutations, which often have small or moderate effects [17]. In particular, most amino-acid changes do not completely destroy a protein’s function, but rather alter its biochemical activity to a greater or lesser extent [18]. The ideal experiment would thus measure the functional effects of many random mutations on many proteins, but such experiments remain challenging [19]. To overcome this experimental limitation, we undertook a computational approach, using biochemically-detailed systems biology models to predict the effects that small perturbations to protein activities will have on the dynamics of the networks in which they function (Fig. 1). We ascribed high and low *dynamical influence* to protein domains for which amino acid substitutions were predicted to have respectively large or small effects on network dynamics. We hypothesized that network dynamics is a synthetic phenotype that is likely subject to natural selection. To test this hypothesis, we compared our predictions of dynamical influence with genomic data on protein domain evolutionary rates in both vertebrates and yeast. We found that dynamical influence is more strongly correlated with evolutionary rate than many previously known correlates. Moreover, dynamical influence remains predictive when knockout phenotype, expression, and network topology are controlled for. Dynamical influence thus offers new insight into selective constraint in protein networks.

**Figure 1.**
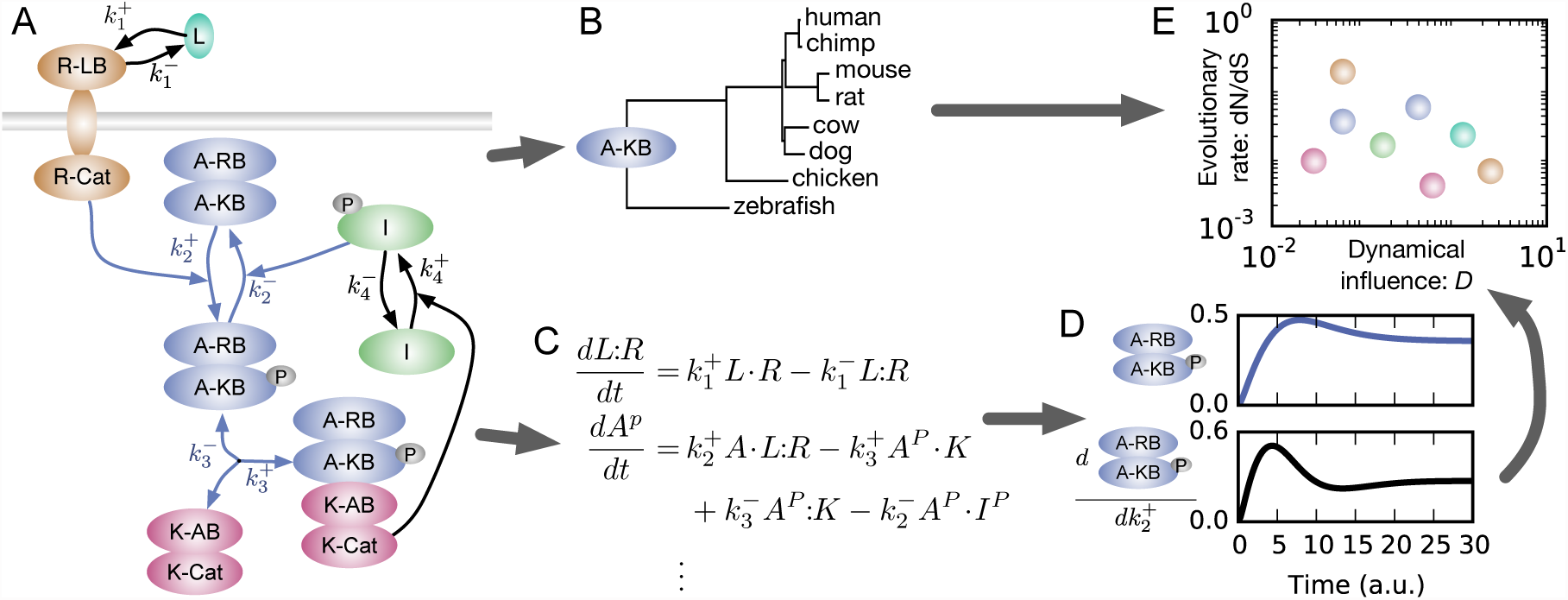
Overview of analysis. A: Illustrative hypothetical signaling network. The dynamical influence of the activator kinase-binding domain (A-KB) is calculated from the influences of the rate constants of the reactions in which it is involved (highlighted in blue): phosphorylation, 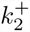; dephos-phorylation, 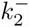; kinase-binding, 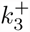; and kinase-unbinding, 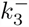. B: Illustrative phylogenetic analysis of the kinase-binding domain of the activator protein. C: Partial list of ordinary differential equations 2 2 2 that model the dynamics of this network. Here all reactions are assumed to be mass-action, but that is not the case in all models analyzed. D: Dynamics of phosphorylated activator protein levels and sensitivity of those dynamics to changes in rate constant 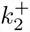, following addition of ligand L. Small increases in 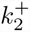 hasten the peak of phosphorylated activator protein and increase its steady-state level. The dynamical influence of rate constant 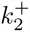 is calculated by summing such sensitivities for all molecular species in the network. E: Illustrative plot comparing dynamical influence and evolutionary rate for all domains in the network. A single multi-domain protein can contribute multiple data points.

## Results and Discussion

### Dynamical influence quantifies the network consequences of small-effect mutations

A biochemically-detailed systems biology model encapsulates vast amounts of molecular biology knowledge in a form that can be used for *in silico* experimentation [20, 21]. In these models, protein biochemical activities are quantified by reaction rate constants *k* [22]. To assess the phenotypic effects of small changes in protein activity caused by mutations, we first calculated the dynamical influence of each reaction rate constant (Materials and Methods). To do so, we calculated how a differential perturbation to that constant would change the concentration time course of each molecular species in the network (Fig. 1D), for biologically-relevant stimuli. We then normalized those changes and integrated the squared changes over time. Lastly, we summed over all molecular species in the network. The dynamical influence of a rate constant is thus the total effect that small changes in that rate constant would have on network dynamics.

The dynamical influence of each reaction rate constant quantifies its importance to network dynamics, but there is little data on evolutionary divergence of reaction rate constants to which we can compare. To compare with the abundant genomic data detailing sequence divergence at the domain level, we aggregated the influences of reaction rate constants for all reactions in which a given protein domain is involved. Whenever possible, we analyzed at the domain level, because that is the level at which distinct functions can be assigned to distinct regions of protein sequence [23]. Thus, we defined the dynamical influence *D* of a domain to be the geometric mean of the dynamical influences of the reaction rate constants for reactions in which it participates (Fig. 1A).

### Dynamical influence is correlated with protein domain evolutionary rate

To test whether dynamical influence is informative about protein evolution, we analyzed dynamic protein network models from BioModels [24], a database which not only collects such models but also annotates them with links to other bioinformatic databases [25, 26]. We considered only models with experimental validation that were formulated in terms of molecular species and reactions, were runnable as ordinary differential equations, and contained at least eight distinct UniProt protein annotations. In total, we studied 12 vertebrate [27, 28, 29, 30, 31, 32, 33, 34, 35, 36, 37, 38] and 6 yeast [39, 40, 41, 42, 43, 44] signaling and biosynthesis models. We further annotated these models to connect molecular species and reactions with particular protein domains (Dataset S1). For each model, we calculated dynamical influences for each reaction rate constant using the stimulation conditions considered in the model’s original publication (Text S1).

Using this novel method, we determined protein domain dynamical influence and evolutionary rate for 18 conserved signaling and metabolic networks (Fig. 2). We quantified the strength of the relationship between dynamical influence and evolutionary rate using Spearman rank correlations (*ρ*), and in 10 of 12 vertebrate networks and 6 of 6 yeast networks, we found a negative correlation. This is consistent with the expectation that most sequences and networks evolve primarily under purifying selection [45], in which natural selection is primarily acting to remove deleterious mutations from the population. Mutations in protein domains with high dynamical influence are predicted to have greater phenotypic effect and thus, in general, be more deleterious. So mutations in those domains are more efficiently removed, and those domains evolve more slowly. Demonstrating the strength of our approach, the two exceptional vertebrate models with a positive correlation, visual signal transaction and interleukin 6 (IL-6) signaling, were recently identified as undergoing network-level adaptation in humans using population genetic data [46]. Positively selected molecular changes in rhodopsin associated with changes in absorption wavelength have been shown to effect dose-response behavior in visual signal transduction [47, 48], suggesting that network-level adaptation may compensate for changes in rhodopsin. As part of the innate immune system, IL-6 and its receptor evolve under strong diversifying selection, so downstream proteins may evolve to maintain signal fidelity. Moreover, viruses are known that directly interfere with proteins downstream of IL-6 [49, 50], potentially driving additional adaptation. Dynamical influence is thus predictive not only about purifying selection but also about adaptive selection.

**Figure 2.**
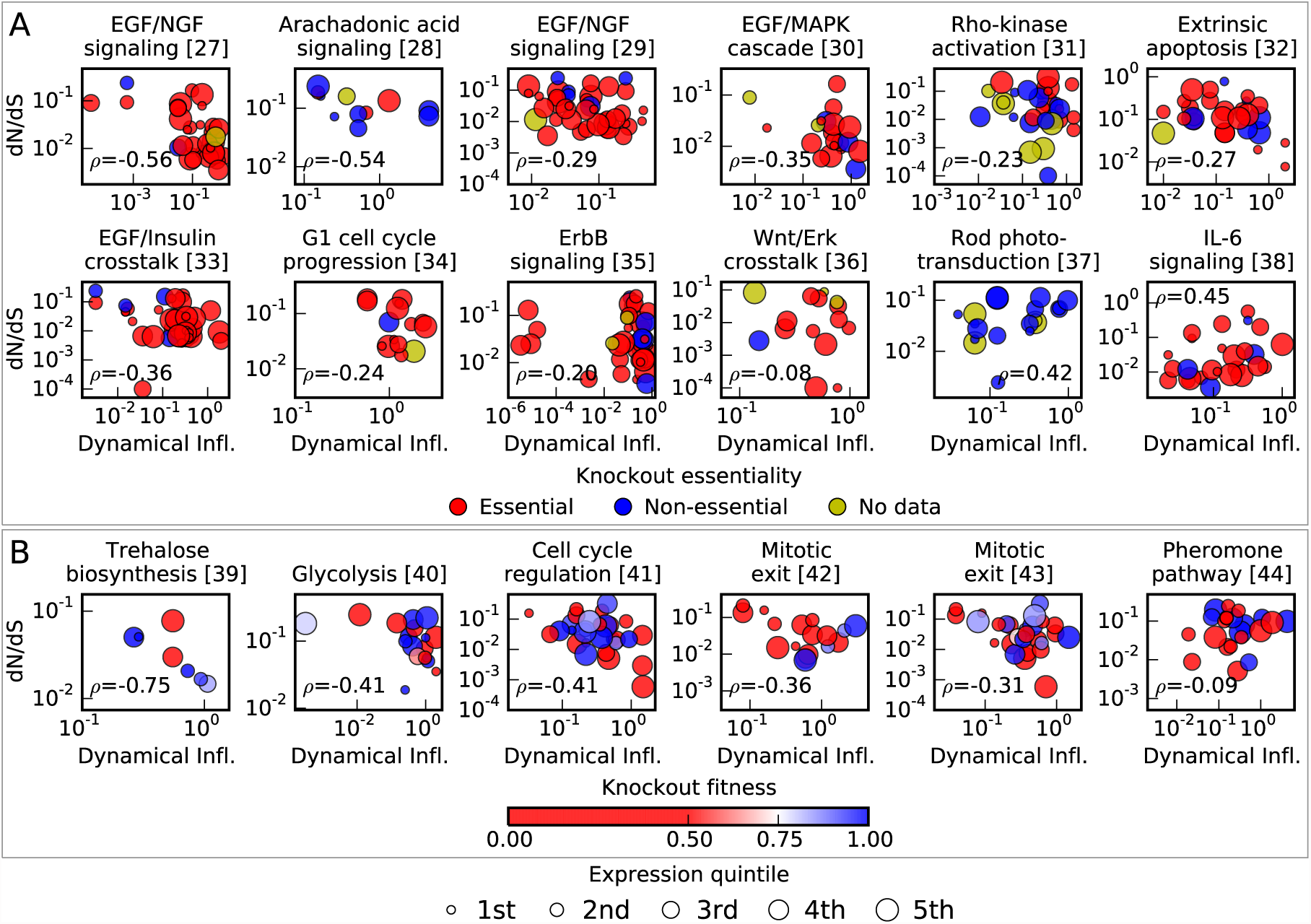
Evolutionary rate is correlated with dynamical influence in signaling and biosyn-thetic networks. Each point represents a protein domain, plotted given its evolutionary rate dN/dS and dynamical influence. Spearman rank correlations *ρ* between dynamical influence and evolutionary rate are generally negative, indicative of widespread purifying selection on network dynamics. Expression level is represented by marker size and is weakly correlated with evolutionary rate but not significantly correlated with dynamical influence (Table 1). A: Vertebrate networks. Knockout essentiality is represented by color, and is not significantly correlated with evolutionary rate or dynamical influence (Table 1). B: Yeast networks. Knockout growth rate is represented by color, with red indicating a more severe phenotype. Knockout growth rate is not significantly correlated with evolutionary rate or dynamical influence (Table 1).

**Table 1:** Overall correlations between evolutionary rate, dynamical influence, and other explanatory variables. Spearman rank (*ρ*) and rank biserial (*rb*) correlation coefficients for variables evolutionary rate dN/dS (*ω*), dynamical influence (*D*), expression breadth (*B*, for vertebrates only), expression level (*X*), interaction degree (*d*), interaction betweenness centrality (*C*), knockout essentiality (*E*), and knockout growth rate (*Gr*, for yeast only). Correlations were combined over all analyzed models using the method of Hunter and Schmidt [51, 52], and p-values were calculated via permutation (Materials and Methods) as well as [53, 54]. Dynamical influence is independently predictive of evolutionary rate, as shown by the negative and statistically significant partial correlation after controlling for all other variables.

|  | correlation | p-value |
| --- | --- | --- |
| $\rho_{\omega,D}$ | -0.23 | <0.0001 |
| $\rho_{\omega,B}$ | -0.25 | 0.0008 |
| $\rho_{\omega,X}$ | -0.14 | 0.0167 |
| $\rho_{\omega,d}$ | -0.21 | 0.0004 |
| $\rho_{\omega,C}$ | -0.18 | 0.0016 |
| $rb_{\omega,E}$ | -0.12 | 0.2061 |
| $\rho_{\omega,Gr}$ | -0.02 | 0.8531 |
| $\rho_{D,B}$ | +0.17 | 0.0090 |
| $\rho_{D,X}$ | +0.09 | 0.0839 |
| $\rho_{D,d}$ | +0.07 | 0.1805 |
| $\rho_{D,C}$ | +0.06 | 0.2794 |
| $rb_{D,E}$ | +0.08 | 0.3378 |
| $\rho_{D,Gr}$ | +0.11 | 0.2429 |
| $\rho_{\omega,D\vert B,X,d,C,E,Gr}$ | -0.13 | 0.0106 |

The strength of the correlation between dynamical influence and protein domain evolutionary rate varies considerably among networks (Tables S1 and S2). To assess the overall strength of the relationship, we combined results across networks as a meta-analysis [51, 52]. This yielded a combined rank correlation of *ρ*_*ω,D*_ = − 0.23, with a permutation p-value of *p <* 10^*-*4^
(Table 1), indicating that the overall pattern is highly unlikely to have arisen by chance.

### Dynamical influence calculation is robust to modeling uncertainties

We measured dynamical influence using hand-built systems biology models; what effect do uncertainties in these models have on our analysis? To be agnostic about what aspects of network dynamics are critical to fitness, in calculating dynamical influence we summed over the integrated sensitivities of all molecular species in the network. It is, however, often evident that the builders of each model had specific molecular species in which they were most interested. If we restricted our dynamical influence calculation to those species (Text S1), we found very similar correlation with domain evolutionary rate (Fig. 3A). Our results are thus not strongly sensitive to which aspects of network function are assumed to be subject to natural selection.

**Figure 3:**
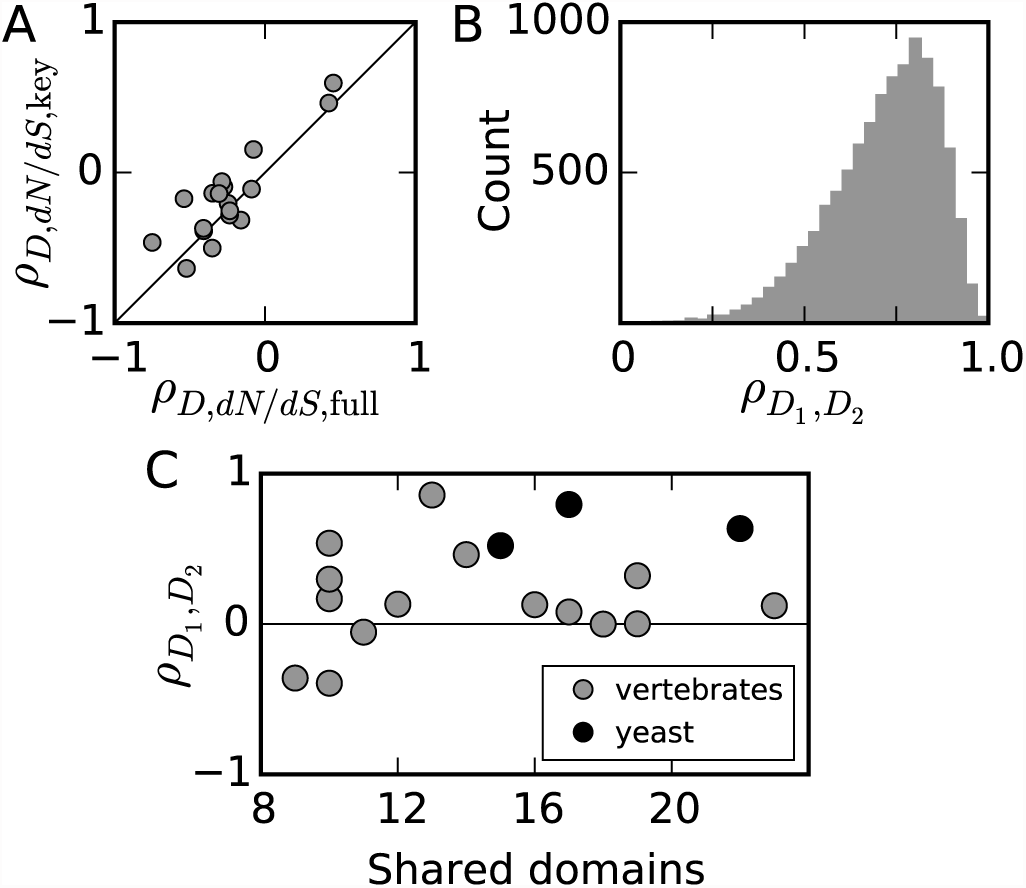
Effects of model uncertainty on dynamical influence. A: The correlation between dynamical influence and evolutionary rate (dN/dS) is similarly strong in all models if dynamical influence is evaluated using all model molecular species (“full”) or only those deemed most important by the authors of the original study (“key”). B: Dynamical influences are strongly correlated between biologically-plausible parameter sets for a model of EGF/NGF signaling [27]. C: Models with overlapping domains produce positively rank-correlated estimates of dynamical influence.

Given a network model, substantial uncertainty can exist about the values of the rate constants *k* [22], because they are difficult to measure directly and are thus often fit to experimental data on network behavior [55, 56]. To assess the importance of this rate constant uncertainty to our results, we used an ensemble of 2000 sets of rate constants [57] that were previously identified as consistent with experimental data for one of our models of EGF/NGF signaling [27]. We calculated the dynamical influence of all protein domains in the network using all these sets of rate constants and compared 10,000 randomly chosen pairs of sets of dynamical influences to each other. Dynamical influences calculated using different plausible rate constant sets are highly correlated (Fig. 3B), with a median rank correlation of 0.74, indicating that rate constant uncertainty does not strongly affect our results.

In addition to parameter uncertainty, different modelers may also make different assumptions when studying the same network regarding forms of interaction, which molecular players to include, or which conditions to consider. We assessed the effect of these assumptions using the models in our data which consider overlapping protein domains. The rank correlation between dynamical influences calculated for the same domains using different models varied considerably and was stronger for pairs of models with larger numbers of overlapping domains (Fig. 3C and Table S3). Combining all these correlations in a meta-analysis as before, we found an overall correlation of 0.26. For comparison, the correlation between different groups measuring gene expression in log-phase growth of budding yeast is roughly 0.62 [58], while for degree in protein-protein interaction data, the correlation is 0.11 [59]. Thus model uncertainty plays a strong but not dominant role in our analysis, and it is comparable to variables that have previously been found to be informative about evolutionary rate.

The existence of overlapping protein domains might inflate statistical significance in our meta-analyses across models. To control for this, we ordered the models by correlation between dynamical influence and evolutionary rate (negative to positive) and recalculated all correlations, removing any domains that had appeared in a previous model (Tables S4 and S5). Meta-analysis of these new correlations yielded similar results (Table S6) to the analyses with all domains (Table 1), so overlapping domains do not strongly affect our conclusions.

### Dynamical influence predicts evolutionary rates independently from previously known factors

Dynamical influence captures the phenotypic effect of small perturbations to protein domain activity, but how does it correlate with previous factors linked to evolutionary rate? In multicellular organisms, proteins that are expressed in more cell types (i.e., have higher expression breadth) evolve more slowly [60], and this is true in the vertebrate networks we study (Table 1). The significant positive correlation between dynamical influence and expression breadth (Table 1) suggests that protein domains with key roles in these networks exert their effects across multiple tissues, providing a functional explanation for the observed correlation between expression breadth and evolutionary rate.

Expanding from expression breadth, the strongest known correlate with protein evolutionary rate is expression level. Proteins with greater expression evolve more slowly in both yeast [61] and vertebrates [62], which may reflect the costs of protein mis-folding [15, 63, 64, 65] or mis-interaction [66]. In our analysis, we find the expected negative correlation between evolutionary rate and expression level (Table 1), but that correlation is notably weaker than that between evolutionary rate and dynamical influence (Table 1). Indeed, dynamical influence is not significantly correlated with expression level (Table 1), indicating that dynamical influence reveals previously unanticipated evolutionary pressures beyond the strongest previously known correlate.

A significant advantage of our approach is that it captures how molecular inputs are integrated into functional phenotypic outcomes that may be selected upon. One aspect of network biology that has been previously considered in determining protein evolution is topology. Specifically, proteins with more interaction partners (i.e., greater degree) [67] or more central locations within networks (greater betweenness centrality) [68] evolve more slowly. Consistent with this previous work, we find that domain evolutionary rate has a significant negative correlation with both protein degree and betweenness centrality (Table 1). But, intriguingly, dynamical influence of protein domains is not significantly correlated with degree or betweenness centrality (Table 1) of the corresponding proteins. Why is the influence of topology not captured in our dynamics-based analysis of evolutionary rate? Network topology is a crude measure of function; networks with the same topology can have different dynamics and thus different functions [69]. Thus, our focus on network dynamics rather than topology provides novel insight into protein domain evolution by directly quantifying system output.

These assessments of dynamical influence relative to known contributors to protein evolution clearly indicate that our approach has uncovered previously unappreciated constraints on protein evolution. Is this new insight sufficient to explain the conundrums raised by knockout experiments? In our data, we found that the correlation between evolutionary rate and knockout measures of function was so weak as to be insignificant (Table 1), consistent with prior work [12, 11]. Strikingly, across the eleven vertebrate networks that include both essential and non-essential proteins and the six yeast networks (for which knockout growth rate data is available), we find no statistical correlation between dynamical influence and essentiality or knockout growth rate (Table 1). Thus, the highly significant correlation between dynamical influence and evolutionary rate (Fig. 2, Table 1) provides a new perspective on the influence of protein function on evolutionary rate.

But, evolutionary rates are complex and likely integrate selection on multiple processes [2, 3, 4, 5]. To assess the power of our approach in comparison with alternative integrative analyses, we used partial correlation analysis [12]. Across all our networks, we find that when expression, network topology, and knockout effect are controlled for, the correlation between protein domain evolutionary rate and dynamical influence remains statistically significant (Table 1). Because the predictive power of dynamical influence cannot be explained by other factors, it provides novel and previously inaccessible insight into evolutionary rates.

## Conclusions

Dynamical systems biology models offer great promise for developing and testing evolutionary hypotheses [21, 70]. Previous topological and flux-balance analysis of networks has offered some insight into protein evolution [71, 72], but dynamical models contribute substantial biological detail not previously captured by these approaches. We have shown that incorporating that detail can explain the previous lack of correlation between protein function and evolutionary rate. While dynamical models have previously been used to predict the phenotypic effects of mutations [73], here we uniquely compare such predictions with genomic data sets to reveal previously unexplored links to evolutionary rate. Given the rapid pace of progress in systems biology modeling [74], the anticipated advances in model scope and validation will provide even more robust data sets to uncover previously unanticipated factors influencing evolutionary processes.

## Materials and Methods

### Dynamical influence

We defined the dynamical influence *κ*_*i*_ of reaction rate constant *k*_*i*_ by

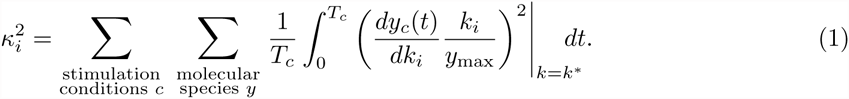

Here *y*_*c*_(*t*) is the time course of molecular species *y* in condition *c*, evaluated using the rate constant values *k*^*\**^ from the original publication. The derivative *dy*_*c*_(*t*)*/dk_i_* of the time course with respect to rate constant *k*_*i*_ measures how sensitive that molecular species is to changes in that rate constant. To make relative comparisons, we normalized these sensitivities by the value *k*_*i*_ of the rate constant and the maximum *y*_max_ of molecular species *y* over all stimulation conditions. To find the total effect of changes in *k*_*i*_, we squared these normalized sensitivities and integrated over the time course of each stimulation condition, and we summed over all molecular species and stimulation conditions.

We defined the dynamical influence *D*_*d*_ of protein domain *d* to be the geometric mean of the influences *κ* of the *N*_*d*_ reaction rate constants for reactions in which that domain participates:

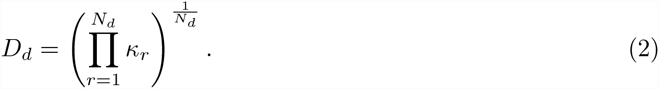

We took a geometric mean because rate constant sensitivities range over orders of magnitude [57].

We downloaded systems biology models in Systems Biology Markup Language (SBML) format [75] from the Feb. 8, 2012 release of BioModels [24]. We calculated dynamical influence for all protein-related biological parameters in each model, using SloppyCell [76] and simulating under the conditions considered in each model’s original paper (Text S1). These parameters represent a variety of biological phenomena, such as binding and catalytic constants and rates of diffusion and production. We considered only those parameters representing rates of biochemical reactions that depend on protein structure, because we expected constraint on those reactions to have the strongest effect on evolutionary rates. Given the dynamical influences *κ* for each reaction constant, we reviewed the literature to determine the protein domain or domains at which the reaction occurs, and we assigned those influences to that domain or domains (Dataset S1).

### Evolutionary rates

We obtained molecular sequences for each protein from Homologene [77] and the Saccharomyces Genome Database [78] (Fig. S1). We quantified protein or domain evolutionary rates using the ratio dN/dS of the rates of nonsynonymous (dN) and synonymous (dS) DNA substitutions, calculated using PAML [79]. Further details and methods for other correlates are in Text S1.

## Acknowledgments

This work was supported by the National Science Foundation, via Graduate Research Fellowship grant DGE-1143953 to BKM. BKM was also supported by an Achievement Rewards for College Scientists scholarship. We thank Tricia Serio for helpful comments on the manuscript.

## 1 Evolutionary rates

UniProt protein ID’s were acquired from the BioModels annotation in the SBML file for each model and converted to NCBI Protein IDs for vertebrates or open reading frame (ORF) numbers for yeast. Some models specified more than one Uniprot ID for a single protein, in cases where there is more than one transcript identified and both appear to perform the same function (for example, MEK1 and MEK2). Where more than one Uniprot ID was specified, we reviewed the model publication and the protein network literature to select a single transcript. In the case of metabolic flux models that track metabolites rather than proteins, we used the names of the enzymes involved in the reactions to find the appropriate protein identifier.

Vertebrate homologous protein alignments were downloaded from the NCBI Homologene database [1], and for each protein in the alignment, nucleotide sequence was downloaded from NCBI Entrez [1]. These nucleotide sequences were then used as a template to back-translate the Homologene protein alignments to nucleotide alignments. Yeast gene information for the 7 species in the tree in Fig. S1B was downloaded from the Saccharomyces Genome Database [2] on Nov. 19, 2012. These gene sequences were translated to protein amino acid sequences using Biopython [3], aligned using ClustalW [4], and then back-translated to aligned nucleotide sequence using the gene sequence as a template.

Protein domain annotation was done manually using literature review, based on information for the human protein in vertebrate models or the *Sa. cerevisiae* protein in yeast. Evolutionary rates were calculated using codeML from PAML Version 4.4b [5], with one dN/dS ratio per tree (model 0), the F3x4 codon substitution model, and a rooted tree, as in [6]. The Mgene=3 setting of codeml was used to estimate a single dN/dS ratio per annotated protein domain. We required a minimum of 4 homologs to include a gene in the analysis, and for each gene any species with more than one homologue was excluded. Because instability is a concern when estimating multiple dN/dS ratios for a single protein sequence, we iterated each codeml run until we acquired three models for which the log-likelihood was within 0.01 of the lowest log-likelihood obtained and then used the model with the lowest log-likelihood.

## 2 Gene expression and specificity

Vertebrate gene expression and tissue specificity data was compiled from the mouse GNF1M dataset [7], downloaded from http://bioGPS.org/downloads. The data consist of microarray probes for a number of tissue types, with each probe’s name including the corresponding gene name, which we mapped to Ensembl gene IDs using Ensembl BioMart [8]. We restricted our analysis to normal adult tissues as in Fig. S2 of [6]. To calculate the expression level corresponding to each microarray probe, we took the arithmetic average over replicates of the same tissue and then took the geometric average over tissues. To calculate the expression level of each gene, we then took the arithmetic average of the probe expression levels corresponding to that gene.

Yeast expression data [9] was obtained from http://younglab.wi.mit.edu/pub/data/orf_transcriptome. txt and used without modification.

## 3 Gene essentiality and dispensibility

We downloaded mouse knockout phenotype data from the Mouse Genome Informatics database [10] at http://www.informatics.jax.org/phenotypes.shtml on July 11, 2011. We assembled phenotype information for homozygous knockouts and coded a gene as essential if it resulted in one of the following phenotypes: abnormal reproductive system physiology, prenatal lethality, perinatal lethality, postnatal lethality, premature death, abnormal reproductive system morphology, lethality at weaning, preweaning lethality, partial lethality, and all sub-phenotypes of these phenotypes. If homozygous knockout of a gene did not cause one or more of these phenotypes we coded it as non-essential. To validate our parsing of this data, we compared against the results of [11].

Data for yeast knockout growth rate on YPD media were obtained from the file Regression Tc1 hom.txt downloaded from the Stanford YDPM database http://www-deletion.stanford.edu/YDPM/YDPM_ index.html on March 13, 2013.

## 4 Network degree and centrality

We downloaded protein-protein interaction data for both humans and yeast from the Interologous Interaction Database [12] on April 20, 2012. These data take the form of a list of interactions between two proteins, and the dataset from which the interaction was curated. Because we were interested in experimentally verified interactions we restricted our analysis to the HPRD, BIND, IntAct, and INNATEDB datasets for humans and the Krogan Core, Yu GoldStd, YeastHigh, YeastLow, and BIND datasets for yeast. We used the python package NetworkX [13] to load these lists of interactions and compute each protein’s degree and its betweenness centrality, which is the fraction of all of the shortest paths between protein pairs in the network that pass through that protein.

## 5 Permutation testing

In our statistical tests, our null model was typically that dynamical influence or evolutionary rate were uncorrelated with other protein domain properties. Because domains share reactions, their influences are not independent, and thus we could not simply permute them to simulate our null model. Instead, we permuted the influences of reaction parameters, which are the most basic unit of our analysis, and then recalculated the influence for each domain based on the new sets of parameter influences. Similarly, many of the correlates with which we compare are defined on the protein level, not the domain level. In these cases, we permuted at the protein level.

## 6 Models considered

### 6.1 EGF/NGF signaling [14]

Brown et al. developed a dynamic model of the EGF/NGF signaling network in rat pheochromocytoma (PC12) cells. We downloaded the SBML file `BIOMD0000000033.xml` from the BioModels database and used it without modification. We simulated this model under two different conditions, 100 ng/ml EGF and 50 ng/ml NGF, as described in [14]. Protein domain, parameter, and reaction assignments are in Dataset S1. In addition to our primary analysis, we calculated dynamical influence over a restricted subset of proteins containing only activated Erk.

### 6.2 Arachodonic acid signaling [15]

In order to gain insights related to anti-inflammatory drug interaction and design, Yang et al. created and studied a dynamic model of the arachadonic acid signaling network in human polymorphonuclear leukocytes. We downloaded the SBML file `BIOMD0000000106.xml` from the BioModels database and used it without modification, integrating the model between 0 and 60 minutes as in [15]. The model contains reactions that create and destroy each tracked molecular species. We excluded these because they are not reactions between modeled proteins. Protein domain, parameter, and reaction assignments are in Dataset S1. In addition to our primary analysis, we calculated dynamical influence over a restricted subset of proteins containing *ω*-LTB4 and 15-Hete.

### 6.3 EGF/NGF signaling [16]

Sasagawa et al. developed a model of ERK signaling networks, with parameters derived by fitting model dynamics to *in vivo* dynamics in PC12 cells, and studied network dynamics under a variety of stimulation conditions. We downloaded SBML file `BIOMD0000000049.xml` from the BioModels database and used it without modification. For all conditions simulated we integrated the network from 0 to 3600 seconds (60 minutes) as in [16]. The SBML file is coded for constant stimulation by 10 ng/ml EGF, and this is the first of the conditions we simulated. We simulated constant stimulation by 10 ng/ml NGF by using SloppyCell to set EGF concentration to 0 and NGF concentration to 10 ng/ml. Sasagawa et al. also investigate the effect of ramping the concentration of EGF (or NGF) from 0 to 1.5 ng/ml over the course of the simulation. To accomplish this we created assignment rules in SloppyCell which updated the EGF (or NGF) concentration at each time step of the integration, setting it equal to 1.5 * (time/3600) ng/ml. As in the fixed simulation conditions, the network was stimulated by EGF or NGF, but not both. Protein domain, parameter, and reaction assignments are in Dataset S1. In addition to our primary analysis, we calculated dynamical influence over a restricted subset of proteins containing only activated Erk.

### 6.4 EGF/MAPK cascade [17]

Schoeberl et al. modeled the EGF signaling pathway, comparing simulated time courses with experimental time courses in HeLa cells under several experimental conditions. We downloaded the SBML file `BIOMD0000000019.xml` from the BioModels database. The model specifies the value of parameter k5 as a piecewise function of another parameter C, and piecewise functions are not supported by SloppyCell, so we removed the piecewise function from the SBML file and used SloppyCell to create two SBML events that replicate it. We simulated the model under three experimental conditions from [17], namely stimulus with 50, 0.5, and 0.25 ng/ml EGF. For all conditions we simulated from 0 to 60 minutes. The model includes receptor internalization reactions which is not modeled mechanistically, and we excluded these reactions from our analysis. Protein domain, parameter, and reaction assignments are in Dataset S1. In addition to our primary analysis, we calculated dynamical influence over a restricted subset of proteins containing only activated Erk.

### 6.5 Myosin phosphorylation [18]

Maeda et al. developed a computational model of thrombin-dependent Rho-kinase activation and myosin light chain phosphorylation in human umbilical vein endothelial cells. We downloaded the SBML file `BIOMD0000000088.xml` for this model from the BioModels database. Some parameter names were duplicated, so we modified the model SBML file to assign unique parameter names and used the published parameter values (see Dataset S1). The only simulation condition we considered was the 0.01*μM* thrombin stimulation published in the SBML file, and we integrated the model from 0 to 3600 seconds as in [18]. We ignored reactions occurring in the extra-cellular compartment, particularly thrombin/thrombin-receptor interactions, and assigned all other reactions to protein domains as outlined in Dataset S1. We did not calculate a sensitivity for the parameter ratio, because it is always multiplied by another parameter V max whose sensitivity we did calculate. SloppyCell interprets rate laws in terms of changes in concentration rather than changes in amount as called for by the SBML specification, so we adjusted reaction stoichiometries by a factor of 1/(compartment size) for reactants in compartment c2, the only compartment in all the models we consider with size not equal to 1. In addition to our primary analysis, we calculated dynamical influence over a restricted subset of proteins containing phosphorylated myosin light chain (pMLC) and phosphorylated myosin phosphatase targeting subunit 1 (pMYPT1).

### 6.6 Extrinsic apoptosis [19]

Albeck et al. developed a model of TRAIL-induced apoptosis and used it to analyze extrinsic apoptosis in HeLa cells. We downloaded the SBML file `BIOMD0000000220.xml` from the BioModels database and used it without modification. We simulated the model for 10 hours under the 50 ng/ml TRAIL stimulus condition encoded in the SBML file. We excluded extra-cellular reactions involving TRAIL, as well as intra-cellular reactions involving DISC and the TRAIL-DISC complex, because the protein components of the DISC are not specified in the model. We also excluded transport across the mitochondrial membrane and binding of proteins to the inner mitochondrial membrane, because these reactions are not mechanistically specified in the model. Protein domain, parameter, and reaction assignments are in Dataset S1. In addition to our primary analysis, we calculated dynamical influence summing over a restricted subset of proteins containing Caspase 3, Caspase 8, cytosolic Smac, and cytosolic Cytochrome C.

### 6.7 EGF/Insulin crosstalk [20]

Borisov et al. developed a model of the Ras/Erk signaling system that incorporates mechanisms of cross-talk between the EGF and Insulin signaling pathways and tested it in HEK293 cells. We down-loaded the SBML file `BIOMD0000000223.xml` from the BioModels database and used it without modification. This SBML model, however, was altered slightly from the originally published model by adding an extra-cellular compartment of size 34. While the model page on the BioModels website says this allows for the use of the original concentrations of EGF we found that this did not create the correct dynamics. Rather than altering the model by changing the size of the extra-cellular compartment we multiplied the desired concentrations of EGF by 34, which produced the correct model dynamics. We simulated the model under four different experimental conditions, 0.01nM or 1 nM EGF with 0 or 100 nM Insulin, as described in [20]. This model contains a reaction in which Akt1 activates mTor via a chain reaction among a number of proteins that are not included in the model. As a result we only applied this reaction’s sensitivity *κ* to the Akt1 kinase domain, but not to mTor. This is the only reaction in the model in which mTor appears, so mTor was excluded from our analysis of this model. Protein domain, parameter, and reaction assignments are in Dataset S1. In addition to our primary analysis, we calculated dynamical influence summing over a restricted subset of proteins containing active (doubly phosphorylated) Erk and phosphorylated Akt.

### 6.8 G1 cell cycle progression [21]

Haberichter et al. constructed a dynamical model of mammalian G1 cell cycle progression in order to simulate cell cycle progression dynamics in proliferating cells continuously exposed to growth factors. We downloaded the SBML file `BIOMD0000000109.xml` from the BioModels database and used it without modification. We simulated the model under the experimental conditions encoded in the SBML file, integrating for 1000 minutes. The model includes reactions that create and destroy proteins, which we excluded from our analysis because they do not involve interaction with any other protein in the model. Protein domain, parameter, and reaction assignments are in Dataset S1. In addition to our primary analysis, we calculated dynamical influence summing over a restricted subset of proteins containing the two activation states, hypo- and hyper-phosphorylated, of retinoblastoma tumor suppressor protein pRb.

### 6.9 ErbB signaling [22]

Birtwistle et al. built a model of ErbB signaling that describes the response of the signaling network to stimulus of all four ErbB receptors with EGF and HRG (heregulin), comparing model dynamics to dynamics in MCF-7 human breast cancer cells. We downloaded SBML file `BIOMD0000000175.xml` from the BioModels database and used it without modification. Experimental conditions in the paper include stimulation with 0 nM, 0.5 nM, and 10 nM EGF and HRG in each possible combination of those three stimuli, for a total of 12 experimental conditions, and we simulate each of these 12 conditions for 2000 seconds. As in other models we excluded receptor internalization reactions that are not mechanistically described in the model. Protein domain, parameter, and reaction assignments are in Dataset S1. In addition to our primary analysis, we calculated dynamical influence summing over a restricted subset of proteins containing active (doubly phosphorylated) Erk and phosphorylated Akt. In particular, we used the normalized active Erk and Akt concentrations described in [22].

### 6.10 Wnt/Erk crosstalk [23]

Kim et al. created a model of the Erk and Wnt pathways to investigate the effect of a positive feedback loop resulting from crosstalk between Wnt and Erk signaling, and they compared model dynamics with experimental results in HEK293 cells. We downloaded the SBML file `BIOMD0000000149.xml` from the BioModels database and used it without modification. This model includes a protein X which is postulated to mediate the feedback between the two pathways; it is transcribed as a result Wnt signaling and activates B-raf in the Erk network. We did not include reactions for transcription and degradation of protein X, because they are not mechanistically specified. We did, however, include the reaction in which protein X activates B-raf, which occurs on the Ras-binding domain (RBD) of B-raf, because binding at the RBD is the mode of activation of B-raf, and the reaction is modeled in the same way as Ras activation of B-raf. We excluded creation and destruction of *β*-catenin as well as degradation of Axin, since they are not mechanistically described in the model. Protein domain, parameter, and reaction assignments are in Dataset S1. In addition to our primary analysis, we calculated dynamical influence summing over a restricted subset of proteins containing active (doubly phosphorylated) Erk and the *β*-catenin/TCF complex.

### 6.11 Rod phototransduction [24]

Dell’Orco et al. developed a model of rod phototransduction specifically aimed at describing the mechanism of light adaptation in rod cells, verifying it by reproducing the results of several previous light adaptation response experiments in mouse rod cells. We downloaded the SBML file `BIOMD0000000326.xml` from the BioModels database and used it with minor modification. We removed two piecewise assignment rules, because SloppyCell does not handle piecewise rules, and replaced them with SBML events. We simulated six flash intensities, replicating those used to create Figure 7 in the publication, by setting the parameter `flash0Mag` to 1.54, 12.5, 45.8, 184, 800, and 2000. The parameter `kP1_rev` represents the rate of dissociation of phosphodiesterase from activated G-alpha molecules, and is set to zero in this model, so we excluded it from our analysis. Protein domain, parameter, and reaction assignments are in Dataset S1. In addition to our primary analysis, we calculated dynamical influence summing over a restricted subset of molecular species containing only cyclic GMP.

### IL-6 signaling [25]

Singh et al. developed a model of IL-6 signaling encompassing the Jak/STAT as well as MAP kinase pathways in human hepatocytes. We downloaded the SBML file `BIOMD0000000151.xml` from the BioModels database and used it without modification. We simulated the experimental conditions encoded in the model, 10 ng/ml IL-6 stimulus and an initial Shp2 concentration of 100 nM. The model includes transcription, translation, and mRNA translocation for the protein SOCS which are not mechanistically detailed, so we excluded them. Protein domain, parameter, and reaction assignments are in Dataset S1. In addition to our primary analysis, we calculated dynamical influence summing over a restricted subset of proteins containing only active STAT3 dimers in the nucleus.

### 6.13 Trehalose biosynthesis [26]

Smallbone et al. created a model of the trehalose biosynthesis pathway in *Sa. cerevisiae*. We downloaded the SBML file `BIOMD0000000380.xml` from the BioModels database and used it without modification. This model reaches a steady state, and the model was validated by comparing the steady state concentrations of the metabolites in the model with experimental results in yeast experiencing heat shock. We simulated the model under the heat shock condition used in the publication, and ran the model for 50000 seconds, at which point all metabolites had reached steady state concentrations. The model creates Clb2 in a reaction that is not mechanistically specified, and we excluded this reaction from our analysis. Protein domain, parameter, and reaction assignments are in Dataset S1. In addition to our primary analysis, we calculated dynamical influence summing over only the metabolite of interest, trehalose-6-phosphate.

### 6.14 Glycolysis [27]

Talser et al. created a model of carbohydrate flux under oxidative stress conditions in *Sa. cerevisiae*. We downloaded the SBML file `BIOMD0000000247.xml` from the BioModels database and used it with substantial modification. The model file in the BioModels database included unfitted parameter values, but model author Markus Ralser generously provided us with the parameter values they obtained by fitting the model to experimental data, and we reparameterized the SBML file accordingly. We simulated the wild-type experimental conditions encoded in the model for 100 minutes. Protein domain, parameter, and reaction assignments are in Dataset S1. In addition to our primary analysis, we calculated dynamical influence summing over the ratio of NADH to NADPH, the quantity of interest.

### 6.15 Cell cycle regulation [28]

Chen et al. developed a model of the cell cycle in *Sa. cerevisiae* in order to investigate the complex mechanisms of cell cycle control. We downloaded the SBML file `BIOMD0000000056.xml` from the BioModels database and used it with substantial modification. This model was constructed with some reactions combined into assignment rules, making it impossible to use SloppyCell to calculate dynamical influence for individual reaction parameters. We separated these assignment rules into individual reactions and added those reactions to the model file, replacing the assignment rules. In order to verify that the modified model was correct we replicated Figures 3 and 6 from the publication. The model contains reactions causing the degradation of various proteins by SCF, which is not included in the model, and these reactions are not mechanistically described, so we excluded them from our analysis. We simulated the wild-type experimental conditions encoded in the paper for 200 minutes as in Figures 3 and 6 of the publication. Protein domain, parameter, and reaction assignments are in Dataset S1. In addition to our primary analysis, we calculated dynamical influence summing over measures of the timing of cell cycle events; MASS, BUD, ORI, and SPN.

### 6.16 Mitotic exit [29]

Queralt et al. developed a model of the initiation of mitotic exit in *Sa. cerevisiae* induced by down-regulation of the phosphatase Cdc50. We downloaded the SBML file `BIOMD0000000409.xml` from the BioModels database and used it without modification. This model contains reactions creating and destroying proteins Clb2, Cdc20, securin, separase, Cdc5, and Cdc15 which are not mechanistically described in the model, and we excluded these reactions from our analysis. We simulated the wild type conditions encoded in the model for 50 minutes, as in Figure 7 of the publication. Protein domain, parameter, and reaction assignments are in Dataset S1. In addition to our primary analysis, we calculated dynamical influence summing over only the concentration of active separase.

### 6.17 Mitotic exit [30]

Vinod et al. created a computational model of mitotic exit in *Sa. cerevisiae* aimed at investigating the role of separase and Cdc14 endocycles. We downloaded the SBML file `BIOMD0000000370.xml` from the BioModels database and used it with substantial modification. This model was constructed using rate rules rather than reactions in SBML. Because SloppyCell our analysis is focused on reactions, we converted the rate rules to reactions, ensuring that the model remained correct by using it to generate Figure 2 from the publication. The model includes reactions creating, destroying, or (de)activating proteins Clb2, Sic1, Cln1, Cdc20, Cdh1, Swi5, Pds1, Esp1, Cdc5, Polo, and MBF which are not mechanistically detailed, and we excluded these reactions from our analysis. We simulated the wild-type experimental conditions encoded in the model for 120 minutes as in Figure 2 of the publication. Protein domain, parameter, and reaction assignments are in Dataset S1. In addition to our primary analysis, we calculated dynamical influence summing over only the concentration of active separase.

### 6.18 Pheromone pathway [31]

Kofahl and Klipp modelled the dynamics of the *Sa. cerevisiae* pheromone pathway. We downloaded the SBML file `BIOMD0000000037.xml` from the BioModels database and used it without modification. The model includes reactions for the destruction of `Ste2` and the export of `Bar` that are not mechanistically detailed, and we excluded them from our analysis. We simulated the wild-type experimental conditions encoded in the model for 30 minutes, as in [31]. Protein domain, parameter, and reaction assignments are in Dataset S1. In addition to our primary analysis, we calculated dynamical influence summing over only the concentration of complexes M and N, which include Far1 and are required for polarized growth and cell cycle arrest respectively.

## 7 Dataset S1

Excel file of data for network parameters and protein domains. Each model corresponds to two sheets. The first sheet contains the reaction parameters, their dynamical influences, and the reactions they correspond to. The second sheet contains the protein and domain data, including assignment of reactions to domains and corresponding references (as PubMed IDs) [32, 33, 34, 35, 36, 37, 38, 39, 40, 41, 42, 43, 44, 45, 46, 47, 48, 49, 50, 51, 52, 53, 54, 55, 56, 57, 58, 59, 60, 61, 62, 63, 64, 65, 66, 67, 68, 69, 70, 71, 72, 73, 74, 75, 76, 77, 78, 79, 80, 81, 82, 83, 84, 85, 86, 87, 88, 89, 90, 91, 92, 93, 94, 95, 96, 97, 98, 99, 100, 101, 102, 103, 104, 105, 106, 107, 108, 109, 110, 111, 112, 113, 114, 115, 116, 117, 118, 119, 120, 121, 122, 123, 124, 125, 126, 127, 128, 129, 130, 131, 132, 133, 134, 135, 136, 137, 138, 139, 140, 141, 142, 143, 29, 144, 145, 146, 147, 148, 149, 150, 151, 152, 153, 154, 155, 156, 157, 158, 159, 160, 161, 162, 163, 164, 165, 166, 167, 168, 169, 170, 171, 172, 173, 174, 175, 176, 177, 178, 179, 180, 181, 182, 183].

Table S1: Correlations in vertebrate models. Spearman rank (*ρ*) and rank biserial (*rb*) correlation coefficients for variables evolutionary rate dN/dS (*ω*), dynamical influence (*D*), expression breadth (*B*), expression level (*X*), interaction degree (*d*), interaction betweenness centrality (*C*), and knock-out essentiality (*E*). Domains with missing values for any correlate were dropped prior to calculating correlations, and *N* represents the number of domains used in the analysis.

---

**Table S1:**
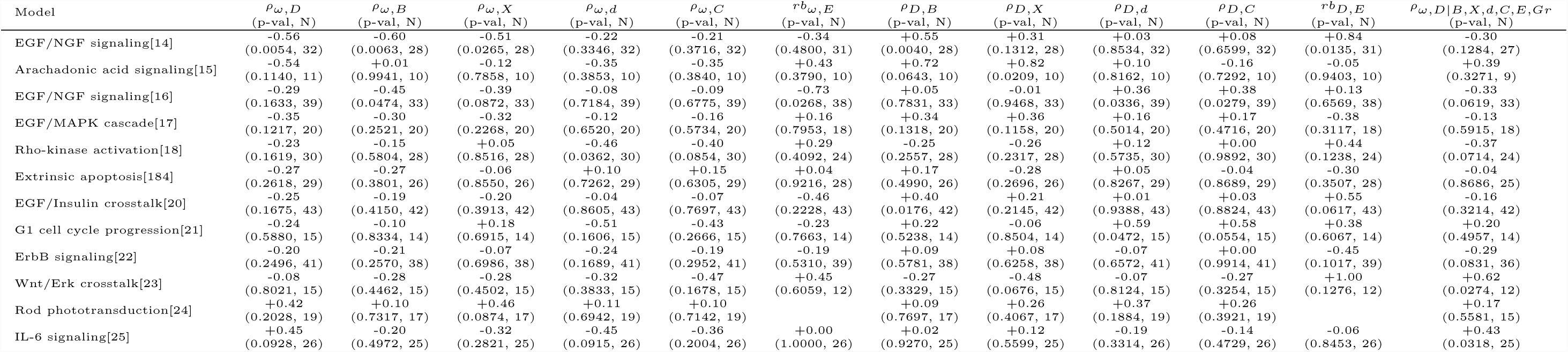
Correlations in vertebrate models. Correlations in yeast models. Spearman rank (*ρ*) and rank biserial (*rb*) correlation coefficients for variables evolutionary rate dN/dS (*ω*), dynamical influence (*D*), expression breadth (*B*), expression level (*X*), interaction degree (*d*), interaction betweenness centrality (*C*), and knock-out essentiality (*E*). Domains with missing values for any correlate were dropped prior to calculating correlations, and N represents the number of domains used in the analysis.

**Table S2:** Correlations in yeast models. Spearman rank (*ρ*) and rank biserial (*rb*) correlation coefficients for variables evolutionary rate dN/dS (*ω*), dynamical influence (*D*), expression level (*X*), interaction degree (*d*), interaction betweenness centrality (*C*), knock-out essentiality (*E*), and knock-out growth rate (*Gr*). Domains with missing values for any correlate were dropped prior to calculating correlations, and *N* represents the number of domains used in the analysis.

| Model | $\rho_{\omega,D}$<br>(p-val, N) | $\rho_{\omega,X}$<br>(p-val, N) | $\rho_{\omega,d}$<br>(p-val, N) | $\rho_{\omega,C}$<br>(p-val, N) | $r^b_{\omega,E}$<br>(p-val, N) | $\rho_{\omega,Gr}$<br>(p-val, N) | $\rho_{D,X}$<br>(p-val, N) | $\rho_{D,d}$<br>(p-val, N) | $\rho_{D,C}$<br>(p-val, N) | $r^b_{D,E}$<br>(p-val, N) | $\rho_{D,Gr}$<br>(p-val, N) | $\rho_{\omega,D\vert B,X,d,C,E,Gr}$<br>(p-val, N) |
| --- | --- | --- | --- | --- | --- | --- | --- | --- | --- | --- | --- | --- |
| Trehalose biosynthesis[26] | -0.75<br>(0.0718, 7) | +0.32<br>(0.5041, 7) | -0.56<br>(0.3728, 5) | -0.90<br>(0.0831, 5) | +0.60<br>(0.2883, 7) | -0.05<br>(0.9161, 7) | -0.14<br>(0.7899, 7) | +0.56<br>(0.3667, 5) | +0.80<br>(0.1370, 5) | -0.20<br>(0.8596, 7) | -0.34<br>(0.4565, 7) | -0.91<br>(0.0502, 5) |
| Glycolysis[27] | -0.41<br>(0.0958, 18) | +0.85<br>(<0.0001, 18) | +0.33<br>(0.1922, 17) | +0.04<br>(0.8707, 17) | +0.12<br>(0.7292, 18) | -0.22<br>(0.3841, 18) | -0.33<br>(0.1815, 18) | -0.20<br>(0.4515, 17) | -0.02<br>(0.9283, 17) | +0.14<br>(0.6558, 18) | -0.03<br>(0.8986, 18) | -0.11<br>(0.6940, 17) |
| Cell cycle regulation[28] | -0.41<br>(0.0207, 29) | -0.32<br>(0.1009, 29) | -0.44<br>(0.0218, 29) | -0.21<br>(0.2923, 29) | -0.08<br>(0.7466, 29) | -0.00<br>(0.9928, 29) | +0.16<br>(0.3959, 29) | +0.07<br>(0.7239, 29) | -0.16<br>(0.3946, 29) | +0.17<br>(0.4428, 29) | -0.14<br>(0.4604, 29) | -0.40<br>(0.0297, 29) |
| Mitotic exit[29] | -0.36<br>(0.2476, 17) | -0.47<br>(0.0549, 17) | -0.42<br>(0.0933, 17) | -0.16<br>(0.5653, 17) | +0.35<br>(0.4030, 17) | -0.26<br>(0.3115, 17) | +0.26<br>(0.2932, 17) | +0.22<br>(0.3782, 17) | +0.02<br>(0.9481, 17) | -0.73<br>(0.0424, 17) | +0.52<br>(0.0247, 17) | +0.04<br>(0.8903, 17) |
| Mitotic exit[30] | -0.31<br>(0.1362, 27) | -0.28<br>(0.2125, 27) | -0.40<br>(0.0646, 27) | -0.16<br>(0.4586, 27) | -0.12<br>(0.6393, 27) | +0.05<br>(0.8377, 27) | +0.25<br>(0.2045, 27) | +0.01<br>(0.9495, 27) | -0.14<br>(0.4863, 27) | -0.08<br>(0.7306, 27) | +0.12<br>(0.5383, 27) | -0.29<br>(0.1464, 27) |
| Pheromone pathway[31] | -0.09<br>(0.6949, 23) | -0.21<br>(0.4093, 23) | -0.17<br>(0.5066, 23) | -0.28<br>(0.2507, 23) | -0.27<br>(0.3980, 23) | +0.23<br>(0.3507, 23) | +0.40<br>(0.0635, 23) | -0.28<br>(0.2048, 23) | +0.16<br>(0.4501, 23) | -0.37<br>(0.1627, 23) | +0.36<br>(0.0987, 23) | -0.21<br>(0.3242, 23) |

**Table S3:**
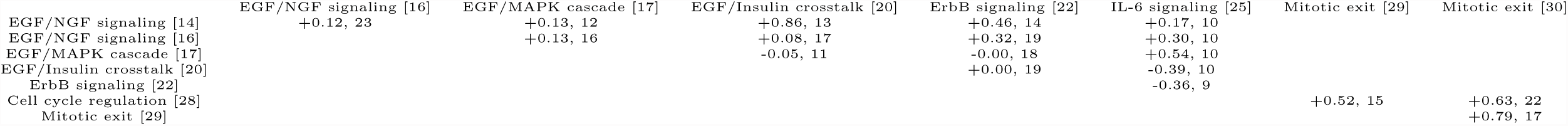
Between-model correlations between protein domain dynamical influences. For each pair of models with at least four overlapping domains, shown is the Spearman rank correlation and number of overlapping domains.

**Table S4:**
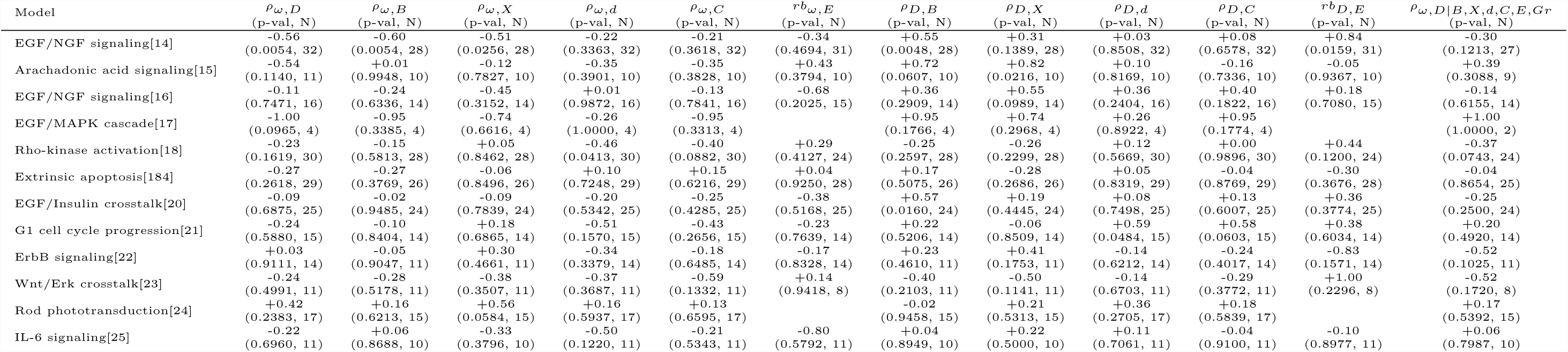
Correlations in yest models with duplicate domains removed. As in Table S1, but removing any domain that appeared in a previous model (in the order shown).

**Table S5:** Correlations in yeast models with overlapping domains removed. As in Table S2, but removing any domain that appeared in a previous model. (in the order shown)

| Model | $\rho_{\omega,D}$<br>(p-val, N) | $\rho_{\omega,X}$<br>(p-val, N) | $\rho_{\omega,d}$<br>(p-val, N) | $\rho_{\omega,C}$<br>(p-val, N) | $r^b_{\omega,E}$<br>(p-val, N) | $\rho_{\omega,Gr}$<br>(p-val, N) | $\rho_{D,X}$<br>(p-val, N) | $\rho_{D,d}$<br>(p-val, N) | $\rho_{D,C}$<br>(p-val, N) | $r^b_{D,E}$<br>(p-val, N) | $\rho_{D,Gr}$<br>(p-val, N) | $\rho_{\omega,D\vert B,X,d,C,E,Gr}$<br>(p-val, N) |
| --- | --- | --- | --- | --- | --- | --- | --- | --- | --- | --- | --- | --- |
| Trehalose biosynthesis[26] | -0.75<br>(0.0718, 7) | +0.32<br>(0.4947, 7) | -0.56<br>(0.3687, 5) | -0.90<br>(0.0828, 5) | +0.60<br>(0.2829, 7) | -0.05<br>(0.9205, 7) | -0.14<br>(0.7788, 7) | +0.56<br>(0.3719, 5) | +0.80<br>(0.1360, 5) | -0.20<br>(0.8569, 7) | -0.34<br>(0.4502, 7) | -0.91<br>(0.0484, 5) |
| Glycolysis[27] | -0.34<br>(0.1970, 16) | +0.85<br>(0.0003, 16) | +0.36<br>(0.1676, 16) | +0.06<br>(0.8342, 16) | +0.13<br>(0.7124, 16) | -0.20<br>(0.4504, 16) | -0.28<br>(0.3008, 16) | -0.23<br>(0.3974, 16) | -0.06<br>(0.8099, 16) | +0.17<br>(0.5935, 16) | -0.11<br>(0.6762, 16) | -0.13<br>(0.6349, 16) |
| Cell cycle regulation[28] | -0.41<br>(0.0207, 29) | -0.32<br>(0.0967, 29) | -0.44<br>(0.0206, 29) | -0.21<br>(0.2930, 29) | -0.08<br>(0.7493, 29) | -0.00<br>(0.9935, 29) | +0.16<br>(0.4067, 29) | +0.07<br>(0.7221, 29) | -0.16<br>(0.3938, 29) | +0.17<br>(0.4467, 29) | -0.14<br>(0.4557, 29) | -0.40<br>(0.0326, 29) |
| Mitotic exit[29] | +1.00<br>(1.0000, 2) |  |  |  |  |  |  |  |  |  |  | +1.00<br>(1.0000, 2) |
| Mitotic exit[30] | +0.50<br>(1.0000, 3) | +0.50<br>(1.0000, 3) | +1.00<br>(0.3316, 3) | +1.00<br>(0.3374, 3) | +0.00<br>(1.0000, 3) | +0.50<br>(1.0000, 3) | -0.50<br>(1.0000, 3) | +0.50<br>(1.0000, 3) | +0.50<br>(1.0000, 3) | +1.00<br>(0.6659, 3) | -0.50<br>(1.0000, 3) | +0.98<br>(0.3313, 3) |
| Pheromone pathway[31] | -0.06<br>(0.7952, 22) | -0.18<br>(0.4792, 22) | -0.05<br>(0.8507, 22) | -0.17<br>(0.4933, 22) | -0.21<br>(0.5026, 22) | +0.19<br>(0.4477, 22) | +0.40<br>(0.0663, 22) | -0.32<br>(0.1402, 22) | +0.15<br>(0.5079, 22) | -0.38<br>(0.1615, 22) | +0.37<br>(0.1014, 22) | -0.18<br>(0.4075, 22) |

**Table S6:** Overall correlations with overlapping domains removed. Meta-analyses as in Table 1, but based on the correlations without overlapping domains (Tables S4 and S5).

|  | correlation | p-value |
| --- | --- | --- |
| $\rho_{\omega,D}$ | -0.22 | 0.0010 |
| $\rho_{\omega,B}$ | -0.20 | 0.0509 |
| $\rho_{\omega,X}$ | -0.05 | 0.5103 |
| $\rho_{\omega,d}$ | -0.19 | 0.0142 |
| $\rho_{\omega,C}$ | -0.18 | 0.0219 |
| $rb_{\omega,E}$ | -0.18 | 0.1499 |
| $\rho_{\omega,Gr}$ | +0.01 | 0.9181 |
| $\rho_{D,B}$ | +0.19 | 0.0300 |
| $\rho_{D,X}$ | +0.07 | 0.3344 |
| $\rho_{D,d}$ | +0.07 | 0.3193 |
| $\rho_{D,C}$ | +0.05 | 0.4546 |
| $rb_{D,E}$ | +0.14 | 0.2057 |
| $\rho_{D,Gr}$ | +0.03 | 0.8096 |
| $\rho_{\omega,D\vert B,X,d,C,E,Gr}$ | -0.20 | 0.0035 |

**Figure S1:**
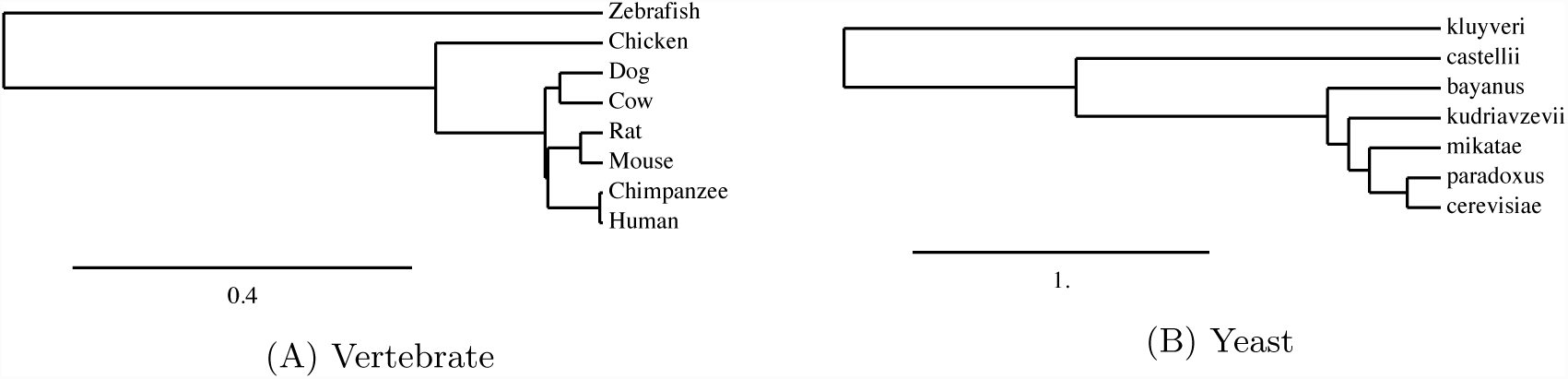
Phylogenetic trees for species used in this study. Branch lengths represent amino acid divergence.

## References

[1] Zuckerkandl E, Pauling L (1965) Evolutionary Divergence and Convergence in Proteins. Evolving Genes and Proteins pp. 97–165.

[2] Pál C, Papp B, Lercher MJ (2006) An integrated view of protein evolution. Nat Rev Genet 7(5):337–48.

[3] Koonin EV, Wolf YI (2006) Evolutionary systems biology: links between gene evolution and function. Curr Opin Biotech 17(5):481–7.

[4] Alvarez-Ponce D (2014) Why Proteins Evolve at Different Rates: The Determinants of Proteins’ Rates of Evolution in Natural Selection: Methods and Applications, ed. Fares MA. (CRC Press), pp. 126–178.

[5] Zhang J, Yang JR (2015) Determinants of the rate of protein sequence evolution. Nature Reviews Genetics 16(7):409–420.

[6] Kimura M, Ota T (1974) On some principles governing molecular evolution. Proc Natl Acad Sci U S A 71(7):2848–52.

[7] Wilson AC, Carlson SS, White TJ (1977) Biochemical evolution. Annu Rev Biochem 46:573–639.

[8] Jordan IK, Rogozin IB, Wolf YI, Koonin EV (2002) Essential genes are more evolutionarily conserved than are nonessential genes in bacteria. Genome Res 12(6):962–968.

[9] Rocha EPC, Danchin A (2004) An analysis of determinants of amino acids substitution rates in bacterial proteins. Mol Biol Evol 21(1):108–16.

[10] Hurst LD, Smith NG (1999) Do essential genes evolve slowly? Curr Biol 9(14):747–50.

[11] Wang Z, Zhang J (2009) Why is the correlation between gene importance and gene evolutionary rate so weak? PLoS Genet 5(1):e1000329.

[12] Liao BY, Scott NM, Zhang J (2006) Impacts of gene essentiality, expression pattern, and gene compactness on the evolutionary rate of mammalian proteins. Mol Biol Evol 23(11):2072–80.

[13] Hirsh AE, Fraser HB (2001) Protein dispensability and rate of evolution. Nature 411(6841):1046–9.

[14] Pál C, Papp B, Hurst LD (2003) Rate of evolution and gene dispensability. Nature 421(January):496–497.

[15] Drummond DA, Bloom JD, Adami C, Wilke CO, Arnold FH (2005) Why highly expressed proteins evolve slowly. Proc Natl Acad Sci U S A 102(40):14338–43.

[16] Hardison RC (2003) Comparative genomics. PLoS Biol 1(2):e58.

[17] Soskine M, Tawfik DS (2010) Mutational effects and the evolution of new protein functions. Nat Rev Genet 11(8):572–82.

[18] Guo HH, Choe J, Loeb LA (2004) Protein tolerance to random amino acid change. Proc Natl Acad Sci U S A 101(25):9205–10.

[19] Fowler DM et al. (2010) High-resolution mapping of protein sequence-function relationships. Nat Methods 7(9):741–6.

[20] Di Ventura B, Lemerle C, Michalodimitrakis K, Serrano L (2006) From in vivo to in silico biology and back. Nature 443(7111):527–33.

[21] Loewe L (2009) A framework for evolutionary systems biology. BMC Syst Biol 3:27.

[22] Gunawardena J (2010) Models in sytems biology: the parameter problem and the meaning of robustness in Elements of Computational Systems Biology, eds. Lodhi HM, Muggleton SH. (John Wiley & Sons, Inc., Hoboken, NJ, USA), pp. 19–47.

[23] Vogel C, Bashton M, Kerrison ND, Chothia C, Teichmann SA (2004) Structure, function and evolution of multidomain proteins. Curr Opin Struct Biol 14(2):208–216.

[24] Le Novère N et al. (2006) BioModels Database: a free, centralized database of curated, published, quantitative kinetic models of biochemical and cellular systems. Nucleic Acids Res 34(Database issue):D689–91.

[25] Li C et al. (2010) BioModels Database: An enhanced, curated and annotated resource for published quantitative kinetic models. BMC Syst Biol 4(1):92.

[26] Chelliah V et al. (2014) BioModels: ten-year anniversary. Nucleic Acids Res 43(November 2014):D542–D548.

[27] Brown KS et al. (2004) The statistical mechanics of complex signaling networks: nerve growth factor signaling. Phys Biol 1(3–4):184–95.

[28] Yang K et al. (2007) Dynamic simulations on the arachidonic acid metabolic network. PLoS Comput Biol 3(3):e55.

[29] Sasagawa S, Ozaki YI, Fujita K, Kuroda S (2005) Prediction and validation of the distinct dynamics of transient and sustained ERK activation. Nat Cell Biol 7(4):365–73.

[30] Schoeberl B, Eichler-Jonsson C, Gilles ED, Müller G (2002) Computational modeling of the dynamics of the MAP kinase cascade activated by surface and internalized EGF receptors. Nat Biotechnol 20(4):370–5.

[31] Maeda A et al. (2006) Ca2+-independent phospholipase A2-dependent sustained Rho-kinase activation exhibits all-or-none response. Genes Cells 11(9):1071–83.

[32] Albeck JG et al. (2008) Quantitative analysis of pathways controlling extrinsic apoptosis in single cells. Mol Cell 30(1):11–25.

[33] Borisov N et al. (2009) Systems-level interactions between insulin-EGF networks amplify mitogenic signaling. Mol Syst Biol 5(256):256.

[34] Haberichter T et al. (2007) A systems biology dynamical model of mammalian G1 cell cycle progression. Mol Syst Biol 3(84):84.

[35] Birtwistle MR et al. (2007) Ligand-dependent responses of the ErbB signaling network: experimental and modeling analyses. Mol Syst Biol 3:144.

[36] Kim D, Rath O, Kolch W, Cho KH (2007) A hidden oncogenic positive feedback loop caused by crosstalk between Wnt and ERK pathways. Oncogene 26(31):4571–9.

[37] Dell’Orco D, Schmidt H, Mariani S, Fanelli F (2009) Network-level analysis of light adaptation in rod cells under normal and altered conditions. Mol Biosyst 5(10):1232–46.

[38] Singh A, Jayaraman A, Hahn J (2006) Modeling regulatory mechanisms in IL-6 signal transduction in hepatocytes. Biotechnol Progr 95(979):850–862.

[39] Smallbone K, Malys N, Messiha HL, Wishart JA, Simeonidis E (2011) Building a kinetic model of trehalose biosynthesis in Saccharomyces cerevisiae. Methods Enzymol 500(ull):355–70.

[40] Ralser M et al. (2007) Dynamic rerouting of the carbohydrate flux is key to counteracting oxidative stress. J Biol 6(4):10.

[41] Chen KKC et al. (2004) Integrative analysis of cell cycle control in budding yeast. Mol Biol Cell 15(8):3841.

[42] Queralt E, Lehane C, Novak B, Uhlmann F (2006) Downregulation of PP2A(Cdc55) phosphatase by separase initiates mitotic exit in budding yeast. Cell 125(4):719–32.

[43] Vinod PK et al. (2011) Computational modelling of mitotic exit in budding yeast: the role of separase and Cdc14 endocycles. J R Soc Interface 8(61):1128–41.

[44] Kofahl B, Klipp E (2004) Modelling the dynamics of the yeast pheromone pathway. Yeast 21(10):831–50.

[45] Haldane JBS (1937) The Effect of Variation of Fitness. Am Nat 71(735):337–349.

[46] Daub JT et al. (2013) Evidence for polygenic adaptation to pathogens in the human genome. Mol Biol Evol 30(7):1544–58.

[47] Imai H et al. (2007) Molecular properties of rhodopsin and rod function. J Biol Chem 282(9):6677– 84.

[48] Yokoyama S, Tada T, Zhang H, Britt L (2008) Elucidation of phenotypic adaptations: Molecular analyses of dim-light vision proteins in vertebrates. Proc Natl Acad Sci U S A 105(36):13480–5.

[49] Chatterjee M, Osborne J, Bestetti G, Chang Y, Moore PS (2002) Viral IL-6-induced cell proliferation and immune evasion of interferon activity. Science 298(5597):1432–5.

[50] Harker JA, Dolgoter A, Zuniga EI (2013) Cell-intrinsic IL-27 and gp130 cytokine receptor signaling regulates virus-specific CD4 T cell responses and viral control during chronic infection. Immunity 39(3):548–59.

[51] Hunter JE, Schmidt FL, Jackson GB (1982) Meta-analysis: cumulating research findings across studies. (Sage Publications, Beverley Hills), p. 175.

[52] Field AP (2001) Meta-analysis of correlation coefficients: a Monte Carlo comparison of fixed- and random-effects methods. Psychol Methods 6(0):161–180.

[53] Freedman D, Lane D (1983) A Nonstochastic Interpretation of Reported Significance Levels. J Bus Econ Stat 1(4):292–298.

[54] Anderson MJ, Legendre P (1999) An empirical comparison of permutation methods for tests of partial regression coeffcients in a linear model. J Stat Comput Simul 62(February 2015):271–303.

[55] Aldridge BB, Burke JM, Lauffenburger DA, Sorger PK (2006) Physicochemical modelling of cell signalling pathways. Nat Cell Biol 8(11):1195–203.

[56] Ashyraliyev M, Fomekong-Nanfack Y, Kaandorp JA, Blom JG (2009) Systems biology: parameter estimation for biochemical models. FEBS J 276(4):886–902.

[57] Gutenkunst RN et al. (2007) Universally sloppy parameter sensitivities in systems biology models. PLoS Comput Biol 3(10):e189.

[58] Csárdi G, Franks A, Choi DS, Airoldi EM (2015) Accounting for Experimental Noise Reveals That mRNA Levels, Amplified by Post-Transcriptional Processes, Largely Determine Steady-State Protein Levels in Yeast. PLoS Genetics 11(5):e1005206.

[59] Plotkin JB, Fraser HB (2007) Assessing the determinants of evolutionary rates in the presence of noise. Mol Biol Evol 24(5):1113–21.

[60] Duret L, Mouchiroud D (2000) Determinants of Substitution Rates in Mammalian Genes: Expression Pattern Affects Selection Intensity but Not Mutation Rate. Mol Biol Evol 17(1):68–70.

[61] Pál C, Papp BB, Hurst LD, Pal C (2001) Highly expressed genes in yeast evolve slowly. Genetics 158(2):927–931.

[62] Subramanian S, Kumar S (2004) Gene expression intensity shapes evolutionary rates of the proteins encoded by the vertebrate genome. Genetics 168(1):373–81.

[63] Drummond DA, Wilke CO (2009) The evolutionary consequences of erroneous protein synthesis. Nat Rev Genet 10(10):715–24.

[64] Yang JR, Zhuang SM, Zhang J (2010) Impact of translational error-induced and error-free mis-folding on the rate of protein evolution. Mol Syst Biol 6(421):421.

[65] Geiler-Samerotte KA et al. (2011) Misfolded proteins impose a dosage-dependent fitness cost and trigger a cytosolic unfolded protein response in yeast. Proc Natl Acad Sci U S A 108:680–685.

[66] Yang JR, Liao BY, Zhuang SM, Zhang J (2012) Protein misinteraction avoidance causes highly expressed proteins to evolve slowly. Proc Natl Acad Sci U S A pp. 831–840.

[67] Fraser HB, Hirsh AE, Steinmetz LM, Scharfe C, Feldman MW (2002) Evolutionary rate in the protein interaction network. Science 296(5568):750–2.

[68] Hahn MW, Kern AD (2005) Comparative genomics of centrality and essentiality in three eukaryotic protein-interaction networks. Mol Biol Evol 22(4):803–6.

[69] Mangan S, Alon U (2003) Structure and function of the feed-forward loop network motif. Proc Natl Acad Sci U S A 100(21):11980–5.

[70] Soyer OS, ed. (2012) Evolutionary Systems Biology. (Springer).

[71] Alvarez-Ponce D, Aguadé M, Rozas J (2011) Comparative genomics of the vertebrate insulin/TOR signal transduction pathway: a network-level analysis of selective pressures. Genome Biol Evol 3:87–101.

[72] Vitkup D, Kharchenko P, Wagner A (2006) Influence of metabolic network structure and function on enzyme evolution. Genome Biol 7(5):R39.

[73] Loewe L, Hillston J (2008) The distribution of mutational effects on fitness in a simple circadian clock. (Springer-Verlag, Berlin), pp. 156–175.

[74] Karr JR et al. (2012) A whole-cell computational model predicts phenotype from genotype. Cell 150(2):389–401.

[75] Hucka M et al. (2003) The systems biology markup language (SBML): a medium for representation and exchange of biochemical network models. Bioinformatics 19(4):524–531.

[76] Myers CR, Gutenkunst RN, Sethna JP (2007) Python unleashed on systems biology. Comput Sci Eng 9(3):34–37.

[77] NCBI Resource Coordinators (2014) Database resources of the National Center for Biotechnology Information. Nucleic Acids Res 42(Database issue):D7–17.

[78] Cherry JM et al. (2012) Saccharomyces Genome Database: the genomics resource of budding yeast. Nucleic Acids Res 40(Database issue):D700–5.

[79] Yang Z (2007) PAML 4: phylogenetic analysis by maximum likelihood. Mol Biol Evol 24(8):1586– 91.

## References

[1] NCBI Resource Coordinators (2014) Database resources of the National Center for Biotechnology Information. Nucleic Acids Res 42(Database issue):D7–17.

[2] Cherry JM et al. (2012) Saccharomyces Genome Database: the genomics resource of budding yeast. Nucleic Acids Res 40(Database issue):D700–5.

[3] Cock PJA et al. (2009) Biopython: freely available Python tools for computational molecular biology and bioinformatics. Bioinformatics 25(11):1422–1423.

[4] Larkin MA et al. (2007) Clustal W and Clustal X version 2.0. Bioinformatics 23(21):2947–2948.

[5] Yang Z (2007) PAML 4: phylogenetic analysis by maximum likelihood. Mol Biol Evol 24(8):1586– 91.

[6] Drummond DA, Wilke CO (2008) Mistranslation-induced protein misfolding as a dominant constraint on coding-sequence evolution. Cell 134(2):341–52.

[7] Su AI et al. (2004) A gene atlas of the mouse and human protein-encoding transcriptomes. Proc Natl Acad Sci U S A 101(16):6062–7.

[8] Kinsella RJ et al. (2011) Ensembl BioMarts: a hub for data retrieval across taxonomic space. Database 2011:bar030.

[9] Holstege FC et al. (1998) Dissecting the Regulatory Circuitry of a Eukaryotic Genome. Cell 95(5):717–728.

[10] Blake JA, Bult CJ, Kadin JA, Richardson JE, Eppig JT (2011) The Mouse Genome Database (MGD): premier model organism resource for mammalian genomics and genetics. Nucleic Acids Res 39(Database issue):D842–8.

[11] Liao BY, Scott NM, Zhang J (2006) Impacts of gene essentiality, expression pattern, and gene compactness on the evolutionary rate of mammalian proteins. Mol Biol Evol 23(11):2072–80.

[12] Brown KR, Jurisica I (2005) Online predicted human interaction database. Bioinformatics 21(9):2076–82.

[13] Hagberg AA, Schult DA, Swart PJ (2008) Exploring network structure, dynamics, and function using {NetworkX}. (Pasadena, CA USA), pp. 11–15.

[14] Brown KS et al. (2004) The statistical mechanics of complex signaling networks: nerve growth factor signaling. Phys Biol 1(3–4):184–95.

[15] Yang K et al. (2007) Dynamic simulations on the arachidonic acid metabolic network. PLoS Comput Biol 3(3):e55.

[16] Sasagawa S, Ozaki YI, Fujita K, Kuroda S (2005) Prediction and validation of the distinct dynamics of transient and sustained ERK activation. Nat Cell Biol 7(4):365–73.

[17] Schoeberl B, Eichler-Jonsson C, Gilles ED, Müller G (2002) Computational modeling of the dynamics of the MAP kinase cascade activated by surface and internalized EGF receptors. Nat Biotechnol 20(4):370–5.

[18] Maeda A et al. (2006) Ca2+-independent phospholipase A2-dependent sustained Rho-kinase activation exhibits all-or-none response. Genes Cells 11(9):1071–83.

[19] Albeck JG et al. (2008) Quantitative analysis of pathways controlling extrinsic apoptosis in single cells. Mol Cell 30(1):11–25.

[20] Borisov N et al. (2009) Systems-level interactions between insulin-EGF networks amplify mitogenic signaling. Mol Syst Biol 5(256):256.

[21] Haberichter T et al. (2007) A systems biology dynamical model of mammalian G1 cell cycle progression. Mol Syst Biol 3(84):84.

[22] Birtwistle MR et al. (2007) Ligand-dependent responses of the ErbB signaling network: experimental and modeling analyses. Mol Syst Biol 3:144.

[23] Kim D, Rath O, Kolch W, Cho KH (2007) A hidden oncogenic positive feedback loop caused by crosstalk between Wnt and ERK pathways. Oncogene 26(31):4571–9.

[24] Dell’Orco D, Schmidt H, Mariani S, Fanelli F (2009) Network-level analysis of light adaptation in rod cells under normal and altered conditions. Mol Biosyst 5(10):1232–46.

[25] Singh A, Jayaraman A, Hahn J (2006) Modeling regulatory mechanisms in IL-6 signal transduction in hepatocytes. Biotechnol Progr 95(979):850–862.

[26] Smallbone K, Malys N, Messiha HL, Wishart JA, Simeonidis E (2011) Building a kinetic model of trehalose biosynthesis in Saccharomyces cerevisiae. Methods Enzymol 500(ull):355–70.

[27] Ralser M et al. (2007) Dynamic rerouting of the carbohydrate flux is key to counteracting oxidative stress. J Biol 6(4):10.

[28] Chen KKC et al. (2004) Integrative analysis of cell cycle control in budding yeast. Mol Biol Cell 15(8):3841.

[29] Queralt E, Lehane C, Novak B, Uhlmann F (2006) Downregulation of PP2A(Cdc55) phosphatase by separase initiates mitotic exit in budding yeast. Cell 125(4):719–32.

[30] Vinod PK et al. (2011) Computational modelling of mitotic exit in budding yeast: the role of separase and Cdc14 endocycles. J R Soc Interface 8(61):1128–41.

[31] Kofahl B, Klipp E (2004) Modelling the dynamics of the yeast pheromone pathway. Yeast 21(10):831–50.

[32] Acehan D et al. (2002) Three-Dimensional Structure of the Apoptosome : Implications for Assembly, Procaspase-9 Binding, and Activation Southwestern Medical Center at Dallas. Structure 9:423–432.

[33] Adams JM, Cory S (2007) The Bcl-2 apoptotic switch in Cancer development and therapy. Oncogene 26(9):1324–37.

[34] Adams MN et al. (2011) Structure, function and pathophysiology of protease activated receptors. Pharmacol Ther 130(3):248–82.

[35] Apanovitch DM, Slep KC, Sigler PB, Dohlman HG (1998) Sst2 is a GTPase-activating protein for Gpa1: purification and characterization of a cognate RGS-Galpha protein pair in yeast. Biochemistry 37(14):4815–22.

[36] Asakawa K, Yoshida S, Otake F, Toh-e A (2001) A novel functional domain of Cdc15 kinase is required for its interaction with Tem1 GTPase in Saccharomyces cerevisiae. Genetics 157(4):1437– 50.

[37] Azzam R et al. (2004) Phosphorylation by cyclin B-Cdk underlies release of mitotic exit activator Cdc14 from the nucleolus. Science 305(5683):516–9.

[38] Barbacci E, Guarino B, Stroh J (1995) The Structural Basis for the Specificity of Epidermal Growth Factor and Heregulin Binding. J Biol Chem.

[39] Barberis M et al. (2005) The yeast cyclin-dependent kinase inhibitor Sic1 and mammalian p27Kip1 are functional homologues with a structurally conserved inhibitory domain. Biochem J 387(Pt 3):639–47.

[40] Batzer AG, Rotin D, Urenã JM, Skolnik EY, Schlessinger J (1994) Hierarchy of binding sites for Grb2 and Shc on the epidermal growth factor receptor. Mol Cell Biol 14(8):5192–201.

[41] Bayascas JR (2010) PDK1: the major transducer of PI 3-kinase actions. Curr Top Microbiol Immunol 346:9–29.

[42] Bhunia A, Mohanram H, Bhattacharjya S (2012) Structural determinants of the specificity of a membrane binding domain of the scaffold protein Ste5 of budding yeast: implications in signaling by the scaffold protein in MAPK pathway. Biochim Biophys Acta 1818(5):1250–60.

[43] Blain SW (2008) Switching Cyclin D-Cdk4 kinase activity on and off. Cell Cycle 7(7):892–898.

[44] Bogner C, Leber B, Andrews DW (2010) Apoptosis: embedded in membranes. Curr Opin Cell Biol 22(6):845–851.

[45] Bos JL, Rehmann H, Wittinghofer A (2007) GEFs and GAPs: critical elements in the control of small G proteins. Cell 129(5):865–77.

[46] Boulares AH et al. (1999) Role of poly(ADP-ribose) polymerase (PARP) cleavage in apoptosis. Caspase 3-resistant PARP mutant increases rates of apoptosis in transfected cells. J Biol Chem 274(33):22932–40.

[47] Brown MT, Cooper JA (1996) Regulation, substrates and functions of src. Biochim Biophys Acta 1287(2-3):121–49.

[48] Burke JR, Deshong AJ, Pelton JG, Rubin SM (2010) Phosphorylation-induced Conformational Changes in the Retinoblastoma Protein Inhibit E2F Transactivation Domain Binding *. J Biol Chem 285(21):16286–16293.

[49] Butty AC, Pryciak PM, Huang LS, Herskowitz I, Peter M (1998) The role of Far1p in linking the heterotrimeric G protein to polarity establishment proteins during yeast mating. Science 282(5393):1511–6.

[50] Calzada A, Sacristán M, Sánchez E, Bueno A (2001) Cdc6 cooperates with Sic1 and Hct1 to inactivate mitotic cyclin-dependent kinases. Nature 412(6844):355–8.

[51] Cameron AJM, Parker PJ (2010) Advances in Enzyme Regulation Protein kinase C A family of protein kinases, allosteric effectors or both ? Adv Enzyme Regul 50(1):169–177.

[52] Camps M (1998) Catalytic Activation of the Phosphatase MKP-3 by ERK2 Mitogen-Activated Protein Kinase. Science 280(5367):1262–1265.

[53] Cheever ML et al. (2008) Crystal structure of the multifunctional Gbeta5-RGS9 complex. Nat Struct Mol Biol 15(2):155–62.

[54] Chen X et al. (1998) Crystal structure of a tyrosine phosphorylated STAT-1 dimer bound to DNA. Cell 93(5):827–39.

[55] Chen Y et al. (2011) A conserved motif within RAP1 has diversified roles in telomere protection and regulation in different organisms. Nat Struct Mol Biol 18(2):213–21.

[56] Chipuk JE, Moldoveanu T, Llambi F, Parsons MJ, Green DR (2010) The BCL-2 family reunion. Mol Cell 37(3):299–310.

[57] Cho JH et al. (2011) Tuning protein autoinhibition by domain destabilization. Nat Struct Mol Biol 18(5):550–5.

[58] Choi Y, Konopka JB (2006) Accessibility of cysteine residues substituted into the cytoplasmic regions of the alpha-factor receptor identifies the intracellular residues that are available for G protein interaction. Biochemistry 45(51):15310–7.

[59] Cote RH (2004) Characteristics of photoreceptor PDE (PDE6): similarities and differences to PDE5. Int J Impot Res 16 Suppl 1:S28–33.

[60] Coughlin SR (2000) Thrombin signalling and protease-activated receptors. Nature 407(6801):258–64.

[61] Dance M, Montagner A, Salles JP, Yart A, Raynal P (2008) The molecular functions of Shp2 in the Ras/Mitogen-activated protein kinase (ERK1/2) pathway. Cell Signal 20(3):453–9.

[62] Daran JM, Dallies N, Thines-Sempoux D, Paquet V, Francois J (1995) Genetic and biochemical characterization of the UGP1 gene encoding the UDP-glucose pyrophosphorylase from Saccharomyces cerevisiae. Eur J Biochem 233(2):520–30.

[63] De Virgilio C et al. (1993) Disruption of TPS2, the gene encoding the 100-kDa subunit of the trehalose-6-phosphate synthase/phosphatase complex in Saccharomyces cerevisiae, causes accumulation of trehalose-6-phosphate and loss of trehalose-6-phosphate phosphatase activity. Eur J Biochem 212(2):315–23.

[64] Delgado ML et al. (2001) The glyceraldehyde-3-phosphate dehydrogenase polypeptides encoded by the Saccharomyces cerevisiae TDH1, TDH2 and TDH3 genes are also cell wall proteins. Microbiology 147(Pt 2):411–7.

[65] Dolan JW, Kirkman C, Fields S (1989) The yeast STE12 protein binds to the DNA sequence mediating pheromone induction. Proc Natl Acad Sci U S A 86(15):5703–7.

[66] Dowell SJ, Bishop AL, Dyos SL, Brown AJ, Whiteway MS (1998) Mapping of a yeast G protein betagamma signaling interaction. Genetics 150(4):1407–17.

[67] Dube P et al. (2005) Localization of the Coactivator Cdh1 and the Cullin Subunit Apc2 in a Cryo-Electron Microscopy Model of Vertebrate APC / C. Mol Cell 20:867–879.

[68] Fallon JL, Halling DB, Hamilton SL, Quiocho FA (2005) Structure of calmodulin bound to the hydrophobic IQ domain of the cardiac Ca(v)1.2 calcium channel. Structure 13(12):1881–6.

[69] Ferguson KM, Lemmon MA, Schlessinger J, Sigler PB (1995) Structure of the high a?nity complex of inositol trisphosphate with a phospholipase C pleckstrin homology domain. Cell 83(6):1037–46.

[70] Ferrezuelo F, Aldea M, Futcher B (2009) Bck2 is a phase-independent activator of cell cycle-regulated genes in yeast. Cell Cycle 8(2):239–52.

[71] Gao C, Chen YG (2010) Dishevelled : The hub of Wnt signaling. Cell Signal 22(5):717–727.

[72] Garrison TR et al. (1999) Feedback phosphorylation of an RGS protein by MAP kinase in yeast. J Biol Chem 274(51):36387–91.

[73] Gartner A et al. (1998) Pheromone-dependent G1 cell cycle arrest requires Far1 phosphorylation, but may not involve inhibition of Cdc28-Cln2 kinase, in vivo. Mol Cell Biol 18(7):3681–91.

[74] George R, Schuller AC, Harris R, Ladbury JE (2008) A phosphorylation-dependent gating mechanism controls the SH2 domain interactions of the Shc adaptor protein. J Mol Biol 377(3):740–7.

[75] Geymonat M, Spanos A, de Bettignies G, Sedgwick SG (2009) Lte1 contributes to Bfa1 localization rather than stimulating nucleotide exchange by Tem1. J Cell Biol 187(4):497–511.

[76] Giorgione JR, Lin JH, McCammon JA, Newton AC (2006) Increased Membrane A?nity of the C1 Domain of Protein Kinase C Compensates for the Lack of Involvement of Its C2 Domain in Membrane Recruitment *. J Biol Chem 281(3):1660–1669.

[77] Goh PY, Lim HH, Surana U (2000) Cdc20 protein contains a destruction-box but, unlike Clb2, its proteolysisis not acutely dependent on the activity of anaphase-promoting complex. Eur J Biochem 267(2):434–49.

[78] Golsteyn RM (2005) Cdk1 and Cdk2 complexes (cyclin dependent kinases) in apoptosis : a role beyond the cell cycle. Cancer Lett 217:129–138.

[79] Good M, Tang G, Singleton J, Reményi A, Lim WA (2009) The Ste5 sca?old directs mating signaling by catalytically unlocking the Fus3 MAP kinase for activation. Cell 136(6):1085–97.

[80] Goto JI, Mikoshiba K (2011) Inositol 1,4,5-Trisphosphate Receptor-Mediated Calcium Release in Purkinje Cells: From Molecular Mechanism to Behavior. Cerebellum.

[81] Gotoh N (2008) Regulation of growth factor signaling by FRS2 family docking/scaffold adaptor proteins. Cancer Sci 99(7):1319–25.

[82] Gottardi CJ, Gumbiner BM (2001) Adhesion signaling : Howβ-catenin interacts with its partners. J Biol Chem pp. 792–794.

[83] Granovsky AE, Rosner MR (2008) Raf kinase inhibitory protein : a signal transduction modulator and metastasis suppressor. Cell Res 18(4):452–457.

[84] Grassie ME, Moffat LD, Walsh MP, Macdonald JA (2011) The myosin phosphatase targeting protein (MYPT) family: A regulated mechanism for achieving substrate specificity of the catalytic subunit of protein phosphatase type 1*δ*. Arch Biochem Biophys 510(2):147–59.

[85] Grauslund M, R onnow B (2000) Carbon source-dependent transcriptional regulation of the mitochondrial glycerol-3-phosphate dehydrogenase gene, GUT2, from Saccharomyces cerevisiae. Can J Microbiol 46(12):1096–100.

[86] Gray CH, Good VM, Tonks NK, Barford D (2003) The structure of the cell cycle protein Cdc14 reveals a proline-directed protein phosphatase. EMBO J 22(14):3524–35.

[87] Griffn JL et al. (2012) Near attack conformers dominate *β*-phosphoglucomutase complexes where geometry and charge distribution reflect those of substrate. Proc Natl Acad Sci U S A 109(18):6910–5.

[88] Guan KL et al. (2000) Negative regulation of the serine/threonine kinase B-Raf by Akt. J Biol Chem 275(35):27354–9.

[89] Gurevich VV, Hanson SM, Song X, Vishnivetskiy SA, Gurevich EV (2011) The functional cycle of visual arrestins in photoreceptor cells. Prog Retin Eye Res 30(6):405–30.

[90] Hanke S, Mann M (2009) The Phosphotyrosine Interactome of the Insulin Receptor Family and Its Substrates. Mol Cell Proteomics pp. 519–534.

[91] Hendrickson C, Meyn MA, Morabito L, Holloway SL (2001) The KEN box regulates Clb2 proteolysis in G1 and at the metaphase-to-anaphase transition. Curr Biol 11(22):1781–7.

[92] Hilioti Z, Chung YS, Mochizuki Y, Hardy CF, Cohen-Fix O (2001) The anaphase inhibitor Pds1 binds to the APC/C-associated protein Cdc20 in a destruction box-dependent manner. Curr Biol 11(17):1347–52.

[93] Hong F et al. (2011) Biochemistry of smooth muscle myosin light chain kinase. Arch Biochem Biophys 510(2):135–146.

[94] Hornig NCD, Knowles PP, McDonald NQ, Uhlmann F (2002) The dual mechanism of separase regulation by securin. Curr Biol 12(12):973–82.

[95] Huang KN, Odinsky SA, Cross FR (1997) Structure-function analysis of the Saccharomyces cerevisiae G1 cyclin Cln2. Mol Cell Biol 17(8):4654–66.

[96] Huang Y, Rich RL, Myszka DG, Wu H (2003) Requirement of both the second and third BIR domains for the relief of X-linked inhibitor of apoptosis protein (XIAP)-mediated caspase inhibition by Smac. J Biol Chem 278(49):49517–22.

[97] Hubbard SR, Till JH (2000) Protein tyrosine kinase structure and function. Annu Rev Biochem 69:373–98.

[98] Hwang LH et al. (1998) Budding yeast Cdc20: a target of the spindle checkpoint. Science 279(5353):1041–4.

[99] Ichiba T et al. (1999) Activation of C3G guanine nucleotide exchange factor for Rap1 by phosphorylation of tyrosine 504. J Biol Chem 274(20):14376–81.

[100] Iyer VR et al. (2001) Genomic binding sites of the yeast cell-cycle transcription factors SBF and MBF. Nature 409(6819):533–8.

[101] Jaspersen SL, Charles JF, Morgan DO (1999) Inhibitory phosphorylation of the APC regulator Hct1 is controlled by the kinase Cdc28 and the phosphatase Cdc14. Curr Biol 9(5):227–36.

[102] Kaburagi Y et al. (1993) Site-directed mutagenesis of the juxtamembrane domain of the human insulin receptor. J Biol Chem 268(22):16610–22.

[103] Kao S, Jaiswal RK, Kolch W, Landreth GE (2001) Identification of the mechanisms regulating the differential activation of the mapk cascade by epidermal growth factor and nerve growth factor in PC12 cells. J Biol Chem 276(21):18169–77.

[104] Kast D, Espinoza-Fonseca LM, Yi C, Thomas DD (2010) Phosphorylation-induced structural changes in smooth muscle myosin regulatory light chain. Proc Natl Acad Sci U S A 107(18):8207– 12.

[105] Kiefer JR et al. (2000) Structural insights into the stereochemistry of the cyclooxygenase reaction. Nature 405(6782):97–101.

[106] Kikuchi A (1999) Roles of Axin in the Wnt Signalling Pathway. Science 11(11):777–788.

[107] Kikuta Y et al. (1998) Purification and Characterization of Recombinant Human Neutrophil Leukotriene B 4-Hydroxylase. Arch Biochem Biophys 355(2):201–205.

[108] Kobashigawa Y et al. (2007) Structural basis for the transforming activity of human Cancer-related signaling adaptor protein CRK. Nat Struct Mol Biol 14(6):503–10.

[109] Komolov KE et al. (2009) Mechanism of rhodopsin kinase regulation by recoverin. J Neurochem 110(1):72–9.

[110] Kraft C, Vodermaier HC, Maurer-Stroh S, Eisenhaber F, Peters JM (2005) The WD40 propeller domain of Cdh1 functions as a destruction box receptor for APC/C substrates. Mol Cell 18(5):543–53.

[111] Kuhn H, Walther M, Kuban RJ (2002) Mammalian arachidonate 15-lipoxygenases Structure, function, and biological implications. Prostaglandins Other Lipid Mediat 69:263–290.

[112] Lee KS, Park JE, Asano S, Park CJ (2005) Yeast polo-like kinases: functionally conserved multitask mitotic regulators. Oncogene 24(2):217–29.

[113] Leeuw T et al. (1998) Interaction of a G-protein beta-subunit with a conserved sequence in Ste20/PAK family protein kinases. Nature 391(6663):191–5.

[114] Lehmann U et al. (2006) Determinants governing the potency of STAT3 activation via the individual STAT3-recruiting motifs of gp130. Cell Signal 18(1):40–9.

[115] Lemmon MA (2009) Ligand-induced ErbB receptor dimerization. Exp Cell Res 315(4):638–48.

[116] Li H, Zhu H, Xu CJ, Yuan J (1998) Cleavage of BID by caspase 8 mediates the mitochondrial damage in the Fas pathway of apoptosis. Cell 94(4):491–501.

[117] Liu F, Hill DE, Cherno? J (1996) Direct binding of the proline-rich region of protein tyrosine phosphatase 1B to the Src homology 3 domain of p130(Cas). J Biol Chem 271(49):31290–5.

[118] Liu HY, Meakin SO (2002) ShcB and ShcC activation by the Trk family of receptor tyrosine kinases. J Biol Chem 277(29):26046–56.

[119] Lou M et al. (2006) The first three domains of the insulin receptor di?er structurally from the insulin-like growth factor 1 receptor in the regions governing ligand specificity. Proc Natl Acad Sci U S A 103(33):12429–34.

[120] MacKay VL et al. (1988) The Saccharomyces cerevisiae BAR1 gene encodes an exported protein with homology to pepsin. Proc Natl Acad Sci U S A 85(1):55–9.

[121] Mashima R, Hishida Y, Tezuka T, Yamanashi Y (2009) The roles of Dok family adapters in immunoreceptor signaling. Immunol Rev 232(1):273–85.

[122] McBride HJ, Yu Y, Stillman DJ (1999) Distinct regions of the Swi5 and Ace2 transcription factors are required for specific gene activation. J Biol Chem 274(30):21029–36.

[123] McDonald CB, Seldeen KL, Deegan BJ, Farooq A (2009) SH3 domains of Grb2 adaptor bind to PXpsiPXR motifs within the Sos1 nucleotide exchange factor in a discriminate manner. Biochemistry 48(19):4074–85.

[124] Mead J et al. (2002) Interactions of the Mcm1 MADS box protein with cofactors that regulate mating in yeast. Mol Cell Biol 22(13):4607–21.

[125] Meyn MA, Melloy PG, Li J, Holloway SL (2002) The destruction box of the cyclin Clb2 binds the anaphase-promoting complex/cyclosome subunit Cdc23. Arch Biochem Biophys 407(2):189–95.

[126] Miller ME, Cross FR, Groeger AL, Jameson KL (2005) Identification of novel and conserved functional and structural elements of the G1 cyclin Cln3 important for interactions with the CDK Cdc28 in Saccharomyces cerevisiae. Yeast 22(13):1021–36.

[127] Miller JJ et al. (2006) Emi1 stably binds and inhibits the anaphase-promoting complex / cyclosome as a pseudosubstrate inhibitor. Genes Dev pp. 2410–2420.

[128] Miller ML et al. (2008) Linear motif atlas for phosphorylation-dependent signaling. Sci Signal 1(35):ra2.

[129] Mittag T et al. (2010) Structure/function implications in a dynamic complex of the intrinsically disordered Sic1 with the Cdc4 subunit of an SCF ubiquitin ligase. Structure 18(4):494–506.

[130] Mori S, Iwaoka R, Eto M, Ohki SY (2009) Solution structure of the inhibitory phosphorylation domain of myosin phosphatase targeting subunit 1. Proteins 77(3):732–5.

[131] Nagao K, Yanagida M (2006) Securin can have a separase cleavage site by substitution mutations in the domain required for stabilization and inhibition of separase. Genes Cells 11(3):247–60.

[132] Nicholson DW (1999) Caspase structure, proteolytic substrates, and function during apoptotic cell death. Cell Death Differ 6(11):1028–42.

[133] Nicholson SE et al. (2000) Suppressor of cytokine signaling-3 preferentially binds to the SHP-2-binding site on the shared cytokine receptor subunit gp130. Proc Natl Acad Sci U S A 97(12):6493–8.

[134] Oberst A et al. (2010) Inducible dimerization and inducible cleavage reveal a requirement for both processes in caspase-8 activation. J Biol Chem 285(22):16632–16642.

[135] Oldham WM, Hamm HE (2008) Heterotrimeric G protein activation by G-protein-coupled receptors. Nat Rev Mol Cell Biol 9(1):60–71.

[136] Ory S, Zhou M, Conrads TP, Veenstra TD, Morrison DK (2003) Protein phosphatase 2A positively regulates Ras signaling by dephosphorylating KSR1 and Raf-1 on critical 14-3-3 binding sites. Curr Biol 13(16):1356–64.

[137] Ow YLP, Green DR, Hao Z, Mak TW (2008) Cytochrome c: functions beyond respiration. Nat Rev Mol Cell Biol 9(7):532–42.

[138] Pacold ME et al. (2000) Crystal structure and functional analysis of Ras binding to its e?ector phosphoinositide 3-kinase gamma. Cell 103(6):931–43.

[139] Pearce LR, Komander D, Alessi DR (2010) The nuts and bolts of AGC protein kinases. Nat Rev Mol Cell Biol 11(1):9–22.

[140] Pearl LH, Barford D (2002) Regulation of protein kinases in insulin, growth factor and Wnt signalling. Curr Opin Struct Biol 12(6):761–7.

[141] Peter M, Gartner A, Horecka J, Ammerer G, Herskowitz I (1993) FAR1 links the signal transduction pathway to the cell cycle machinery in yeast. Cell 73(4):747–60.

[142] Pinheiro AS, Marsh JA, Forman-Kay JD, Peti W (2011) Structural signature of the MYPT1-PP1 interaction. J Am Chem Soc 133(1):73–80.

[143] Queralt E, Igual JC (2004) Functional distinction between Cln1p and Cln2p cyclins in the control of the Saccharomyces cerevisiae mitotic cycle. Genetics 168(1):129–40.

[144] Raaijmakers JH, Bos JL (2009) Specificity in Ras and Rap signaling. J Biol Chem 284(17):10995– 9.

[145] Roskoski R (2010) RAF protein-serine/threonine kinases: structure and regulation. Biochem Biophys Res Commun 399(3):313–7.

[146] Rudberg PC, Tholander F, Andberg M, Thunnissen MMGM, Haeggstro JZ (2004) Leukotriene A 4 Hydrolase IDENTIFICATION OF A COMMON CARBOXYLATE RECOGNITION SITE FOR THE EPOXIDE HYDROLASE AND AMINOPEPTIDASE SUBSTRATES. Biochemistry 279(26):27376–27382.

[147] Savaskan NE, Ufer C, Kühn H, Borchert A (2007) Molecular biology of glutathione peroxidase 4: from genomic structure to developmental expression and neural function. Biol Chem 388(10):1007–17.

[148] Schreiber A et al. (2011) Structural basis for the subunit assembly of the anaphase-promoting complex. Nature 470(7333):227–32.

[149] Shen RF, Tai HH (1998) Thromboxanes: synthase and receptors. J Biomed Sci 5(3):153–72.

[150] Shinohara H, Yasuda T, Yamanashi Y (2004) Dok-1 tyrosine residues at 336 and 340 are essential for the negative regulation of Ras-Erk signalling, but dispensable for. Genes Cells 2:601– 607.

[151] Shirayama M, Zachariae W, Ciosk R, Nasmyth K (1998) The Polo-like kinase Cdc5p and the WD-repeat protein Cdc20p/fizzy are regulators and substrates of the anaphase promoting complex in Saccharomyces cerevisiae. EMBO J 17(5):1336–49.

[152] Shuai K, Liu B (2003) Regulation of JAK-STAT signalling in the immune system. Nat Rev Immunol 3(11):900–11.

[153] Simister PC, Feller SM (2012) Order and disorder in large multi-site docking proteins of the Gab family–implications for signalling complex formation and inhibitor design strategies. Mol Biosyst 8(1):33–46.

[154] Singh P, Wang B, Maeda T, Palczewski K, Tesmer JJG (2008) Structures of rhodopsin kinase in Different ligand states reveal key elements involved in G protein-coupled receptor kinase activation. J Biol Chem 283(20):14053–62.

[155] Smith SO (2010) Structure and activation of the visual pigment rhodopsin. Annu Rev Biophys 39:309–28.

[156] Smrcka AV (2008) G protein betagamma subunits: central mediators of G protein-coupled receptor signaling. Cell Mol Life Sci 65(14):2191–214.

[157] Song MS, Salmena L, Pandolfi PP (2012) The functions and regulation of the PTEN tumour suppressor. Nat Rev Mol Cell Biol 13(5):283–296.

[158] Staal FJT, Clevers HC (2003) Wnt signaling in the thymus. Curr Opin Immunol 15(2):204–8.

[159] Stefan MI, Edelstein SJ, Le Novère N (2008) An allosteric model of calmodulin explains Differential activation of PP2B and CaMKII. Proc Natl Acad Sci U S A 105(31):10768–73.

[160] Stennicke HR et al. (1999) Caspase-9 can be activated without proteolytic processing. J Biol Chem 274(13):8359–62.

[161] Stratton HF, Zhou J, Reed SI, Stone DE (1996) The mating-specific G(alpha) protein of Saccharomyces cerevisiae downregulates the mating signal by a mechanism that is dependent on pheromone and independent of G(beta)(gamma) sequestration. Mol Cell Biol 16(11):6325–37.

[162] Suzuki Y, Nakabayashi Y, Takahashi R (2001) Ubiquitin-protein ligase activity of X-linked inhibitor of apoptosis protein promotes proteasomal degradation of caspase-3 and enhances its anti-apoptotic effect in Fas-induced cell death. Proc Natl Acad Sci U S A 98(15):8662–7.

[163] Swift S et al. (2006) Role of the PAR1 receptor 8th helix in signaling: the 7-8-1 receptor activation mechanism. J Biol Chem 281(7):4109–16.

[164] Traverso EE et al. (2001) Characterization of the Net1 cell cycle-dependent regulator of the Cdc14 phosphatase from budding yeast. J Biol Chem 276(24):21924–31.

[165] Uhlmann F, Wernic D, Poupart MA, Koonin EV, Nasmyth K (2000) Cleavage of cohesin by the CD clan protease separin triggers anaphase in yeast. Cell 103(3):375–86.

[166] Vadas O, Burke JE, Zhang X, Berndt A, Williams RL (2011) Structural basis for activation and inhibition of class I phosphoinositide 3-kinases. Sci Signal 4(195):re2.

[167] Varghese JN et al. (2002) Structure of the extracellular domains of the human interleukin-6 receptor alpha-chain. Proc Natl Acad Sci U S A 99(25):15959–64.

[168] Viadiu H, Stemmann O, Kirschner MW, Walz T (2005) Domain structure of separase and its binding to securin as determined by EM. Nat Struct Mol Biol 12(6):552–3.

[169] Wall MA et al. (1995) The Structure of the G Protein Heterotrimer GialPlyz. Cell 83:1047–1058.

[170] Ward CW, Lawrence MC, Streltsov Va, Adams TE, McKern NM (2007) The insulin and EGF receptor structures: new insights into ligand-induced receptor activation. Trends Biochem Sci 32(3):129–37.

[171] Wäsch R, Cross FR (2002) APC-dependent proteolysis of the mitotic cyclin Clb2 is essential for mitotic exit. Nature 418(6897):556–62.

[172] Waters SB et al. (1995) Desensitization of Ras activation by a feedback disassociation of the SOS-Grb2 complex. J Biol Chem 270(36):20883–6.

[173] Wehrman T et al. (2007) Structural and mechanistic insights into nerve growth factor interactions with the TrkA and p75 receptors. Neuron 53(1):25–38.

[174] Weinreich M, Liang C, Chen HH, Stillman B (2001) Binding of cyclin-dependent kinases to ORC and Cdc6p regulates the chromosome replication cycle. Proc Natl Acad Sci U S A 98(20):11211–7.

[175] Wilmes GM et al. (2004) Interaction of the S-phase cyclin Clb5 with an “RXL” docking sequence in the initiator protein Orc6 provides an origin-localized replication control switch. Genes Dev 18(9):981–91.

[176] Winters MJ, Pryciak PM (2005) Interaction with the SH3 domain protein Bem1 regulates signaling by the Saccharomyces cerevisiae p21-activated kinase Ste20. Mol Cell Biol 25(6):2177–90.

[177] Wolgemuth DJ (2008) Function of cyclins in regulating the mitotic and meiotic cell cycles in male germ cells. Cell Cycle 7:3509–3513.

[178] Won AP, Garbarino JE, Lim WA (2011) Recruitment interactions can override catalytic interactions in determining the functional identity of a protein kinase. Proc Natl Acad Sci U S A 108(24):9809–14.

[179] Xiao B et al. (2003) Crystal structure of the retinoblastoma tumor suppressor protein bound to E2F and the. Proc Natl Acad Sci U S A 100(5):2363–2368.

[180] Yamada T, Komoto J, Watanabe K, Ohmiya Y, Takusagawa F (2005) Crystal Structure and Possible Catalytic Mechanism of Microsomal Prostaglandin E Synthase Type 2 (mPGES-2). Structure 2:1163–1176.

[181] Zähringer H, Burgert M, Holzer H, Nwaka S (1997) Neutral trehalase Nth1p of Saccharomyces cerevisiae encoded by the NTH1 gene is a multiple stress responsive protein. FEBS Lett 412(3):615–20.

[182] Zoncu R, Efeyan A, Sabatini DM (2011) mTOR: from growth signal integration to Cancer, diabetes and ageing. Nat Rev Mol Cell Biol 12(1):21–35.

[183] van Es JH et al. (1999) Identification of APC2, a homologue of the adenomatous polyposis coli tumour suppressor. Curr Biol 9(2):105–8.

[184] Albeck JG, Burke JM, Spencer SL, Lauffenburger DA, Sorger PK (2008) Modeling a snap-action, variable-delay switch controlling extrinsic cell death. PLoS Biol 6(12):2831–52.

